# Epigenetic coordination of transcriptional and translational programs in hypoxia

**DOI:** 10.1101/2023.09.16.558067

**Authors:** Kathleen Watt, Bianca Dauber, Krzysztof J. Szkop, Laura Lee, Predrag Jovanovic, Shan Chen, Ranveer Palia, Julia A. Vassalakis, Tyler T. Cooper, David Papadopoli, Laìa Masvidal, Michael Jewer, Kristofferson Tandoc, Hannah Plummer, Gilles A. Lajoie, Ivan Topisirovic, Ola Larsson, Lynne-Marie Postovit

**Affiliations:** Department of Oncology-Pathology, Science for Life Laboratory, Karolinska Institutet, Stockholm, Sweden; Department of Biomedical and Molecular Sciences, Queen’s University, Kingston, ON, Canada; Department of Oncology, University of Alberta, Edmonton, AB, Canada; Lady Davis Institute, Sir Mortimer B. Davis Jewish General Hospital, Montréal, QC, Canada; Division of Experimental Medicine, McGill University, Montréal, QC, Canada; Department of Biochemistry, Western University, London, ON, Canada; Gerald Bronfman Department of Oncology and Department of Biochemistry, McGill University, Montréal, QC, Canada

## Abstract

Adaptation to cellular stresses entails an incompletely understood coordination of transcriptional and post-transcriptional gene expression programs. Here, we quantified hypoxia-dependent transcriptomes, epigenomes, and translatomes in T47D breast cancer cells and H9 human embryonic stem cells. This revealed pervasive changes in transcription start site (TSS) selection associated with nucleosome repositioning and alterations in H3K4me3 distribution. Importantly, hypoxia-associated TSS switching was induced or reversed via pharmacological modulation of H3K4me3 in the absence of hypoxia, defining a role for H3K4me3 in TSS selection independent of HIF1-transcriptional programs. By remodelling 5’UTRs, TSS switching selectively alters protein synthesis, including enhanced translation of mRNAs encoding pyruvate dehydrogenase kinase 1 (*PDK1)* that is essential for the metabolic adaptation to hypoxia. These results demonstrate a previously unappreciated mechanism of translational regulation during hypoxia driven by epigenetic reprogramming of the 5’UTRome.

## INTRODUCTION

Cellular plasticity underlies adaptive responses to microenvironment alterations, and is fundamental to cancer cell survival and metastasis^1^. Plasticity requires coordinated reprogramming of gene expression at transcriptional and post-transcriptional levels to reshape the proteome^2^. Cancer cells commonly adapt to hypoxia, which can enhance plasticity and enrich for cancer stem-cell-like phenotypes^1,3^. Hypoxia imposes metabolic restrictions impacting gene expression^4,5^ and the epigenome^6–8^. This includes accumulation of trimethylation of lysine 4 of the histone H3 subunit (H3K4me3) that occurs around transcription start sites (TSSs) and correlates with the degree and consistency of transcriptional activation^9–12^. H3K4 methylation is deposited by Complex Proteins Associated with Set1 (COMPASS) methyltransferase complexes and is erased by Jumonji C (JMJC) histone demethylases, including histone lysine demethylase 5 (KDM5)^13^. Hypoxia-associated HIF1 stabilization promotes transcription of several JMJC histone demethylases^14–16^, which is counteracted by a reduction in their activity when O_2_ is limiting^17^. This leads to accumulation of H3K4 methylation under hypoxia, remodelling of chromatin, and enhanced cellular plasticity^6,8^. However, the effect of this oxygen-sensing capacity of chromatin on processes such as TSS selection is still elusive.

Cap-dependent mRNA translation is suppressed under hypoxia^2,5^ via inhibition of the mechanistic Target of Rapamycin (mTOR)^18^ and subsequent reduction in eIF4F levels^19^. Hypoxia also induces the integrated stress response (ISR), wherein eIF2α phosphorylation-dependent suppression of eIF2B attenuates ternary complex recycling and initiator tRNA delivery^5,20^. This reprogramming of the translational apparatus in hypoxia reduces global protein synthesis while allowing selective translation of mRNAs encoding central regulators of stress responses^5^ that contain unique 5’UTR features. For example, mRNAs harbouring upstream open reading frames (uORFs)^21^, including ATF4, are selectively translated under hypoxia^22^. Other 5’UTR features, such as length^23^ and the presence of 5’terminal oligopyrimidine (TOP) motifs render translation selectively mTOR-sensitive^23–25^. Finally, mRNAs encoding stem cell-associated factors (NODAL, SNAIL, NANOG) encompass 5’UTR isoforms that contain features that enhance translation under hypoxia^26^. These findings suggest that in addition to reprogramming of the translational machinery, changes in the composition of 5’UTRs may also drive adaptive protein synthesis under hypoxia.

Here, by profiling epigenomes, transcriptomes, 5’UTRomes and translatomes under normoxia and hypoxia, we uncovered a hitherto unappreciated gene expression mechanism wherein epigenomic alterations facilitate hypoxia-induced changes in mRNA translation. We show that perturbations in H3K4me3 induce abundant alterations in TSS selection leading to a gain and/or loss of 5’UTR elements which, in parallel with hypoxia-induced alterations of the protein synthesis machinery, contributes to an adaptive translational response. This mechanism of TSS switching regulates synthesis of key factors that allow metabolic adaptations to hypoxia, including pyruvate dehydrogenase kinase 1 (PDK1). Many 5’UTR isoform changes occur independent of altered transcript abundance and appear to depend on reduced KDM5 activity leading to expansion and redistribution of H3K4me3. Indeed, pharmacological inhibition of KDM5 mimics hypoxia-induced TSS switching and modulates the proteome in the absence of HIF1 stabilization or alterations in mRNA abundance. Conversely, inhibition of MLL-containing COMPASS methyltransferases blocked a subset of TSS switching and reduced cellular proliferation under hypoxia. Collectively, hypoxia-induced changes in H3K4me3 alter TSS selection to establish an adaptive translatome.

## RESULTS

### Hypoxia-induced TSS switching results in extensive remodeling of 5’UTRs

We previously observed that several stem cell-associated factors have multiple 5’UTR isoforms, with some preferentially translated under hypoxia^26^. Divergent cell types respond to hypoxia with differing kinetics, modifying gene expression and regulators of mRNA translation at different time points and oxygen concentrations. Accordingly, we explored hypoxia-associated 5’UTR isoform dynamics by performing nanoCAGE sequencing on total mRNAs from T47D breast cancer cells and H9 human embryonic stem cells (hESCs) grown for 48 or 24 hrs, respectively, in hypoxia (0.5% O_2_) or normoxia (20% O_2_) (**Fig 1a**). These conditions were chosen as they were previously shown to induce *LOX1* mRNA and HIF1α protein levels, suppress mTOR signalling, and induce ISR^26–28^. Approximately 20,000 RefSeq transcripts were detected at a sequencing depth approaching saturation (**Fig S1a-b**). Both cell lines showed increased expression of hypoxia-associated transcripts and condition-dependent separation of samples in a principal component analysis (**Fig S1c-d**). Consistent with previous observations^23^, the weighted mean 5’UTR length (for expressed 5’UTR isoforms) was often shorter than indicated in the RefSeq database (**Fig S1e**). Using reproducible nanoCAGE reads, we defined TSS clusters for each transcript (see methods) representing distinct 5’UTR isoforms. In total, we detected > 70,000 TSS clusters in both T47D and H9 cells and found that > 80% of protein-coding transcripts manifest more than one 5’UTR isoform (**Fig 1b**). Moreover, 5’UTR availability was modulated in > 20% of detected protein-coding transcripts in T47D and H9 cells after excluding TSS switching impacting ORFs (**Fig 1c, Fig S1f, Supplemental File S1**). Interestingly, TSS switching events occurred in transcripts where expression was increased, decreased, or unchanged in response to hypoxia (**Fig 1d**) (**Supplemental File S2**), thus suggesting that TSS alterations are independent of changes in overall transcription. This is exemplified by SH3BP2, whose transcript levels increased under hypoxia alongside a switch in 5’UTR isoform expression (**Fig 1e**), and PELP1, where TSS clusters were gained or lost under hypoxia in the absence of changes in overall mRNA level (**Fig 1f**).

**Figure 1:**
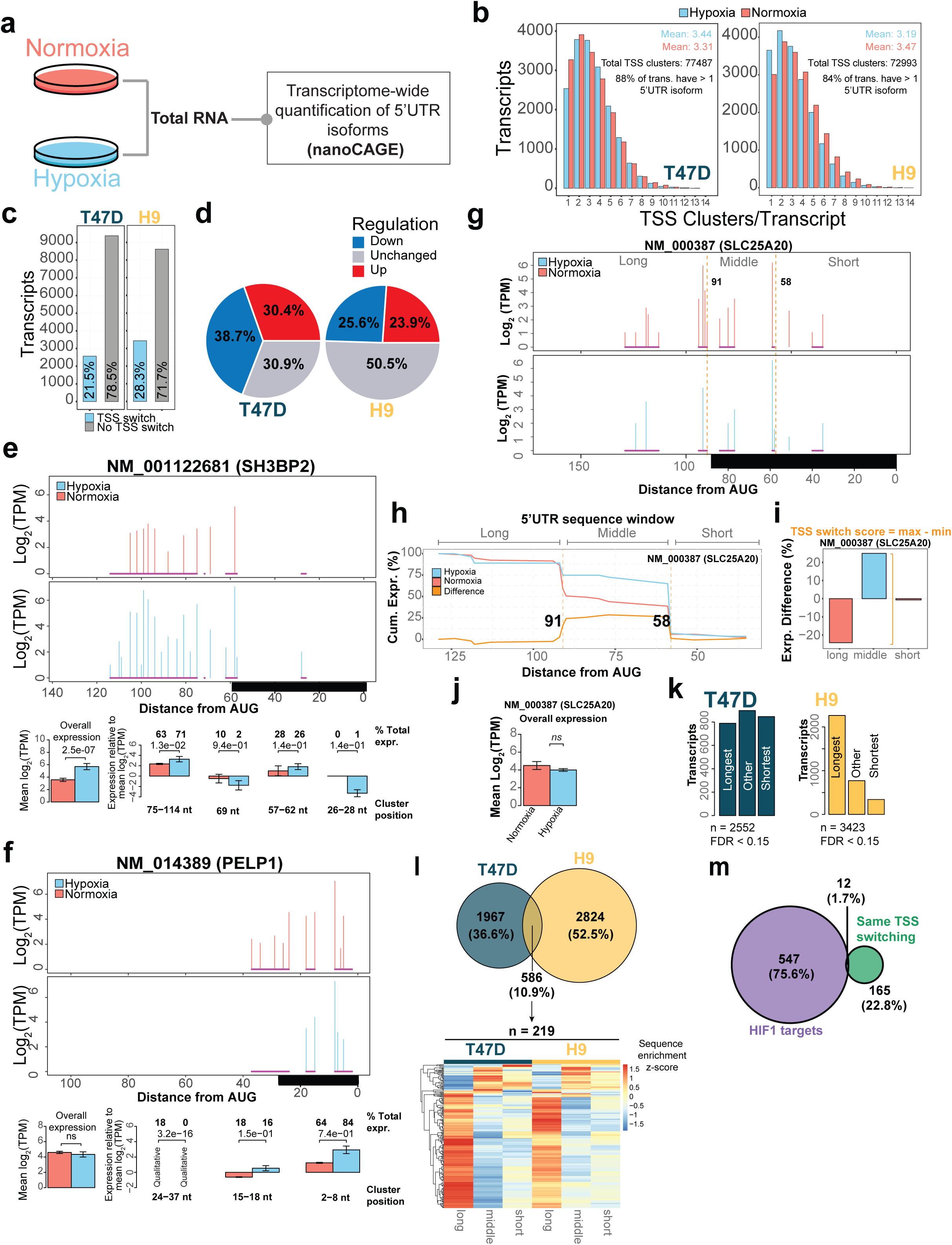
Hypoxia-induced TSS switching results in extensive remodeling of 5’UTRs. **a)** T47D (n = 3 hypoxia, n = 2 normoxia), and H9 (n = 4) cells were cultured under hypoxia (0.5% O_2_) or normoxia (20% O_2_). Total RNA was isolated and transcriptome-wide quantification of 5’UTR isoforms was carried out using nanoCAGE sequencing. **b)** Histograms showing the number of TSS clusters per protein-coding transcript in hypoxia and normoxia in T47D (left) and H9 (right) cells. **c)** Barplot indicating the proportions of protein-coding transcripts with altered TSS usage between hypoxia and normoxia (FDR < 0.15) in T47D (n = 2,552), and H9 cells (n = 3,423). **d)** Pie charts indicating the proportions of transcripts with TSS switching from **c** that have significantly (FDR < 0.15) increased (up), decreased (down), or unchanged overall mRNA expression between hypoxia and normoxia in T47D and H9 cells. Expression was measured using RNA sequencing and differentially expressed genes were identified using anota2seq. **e)** An example of a transcript that undergoes a quantitative hypoxia-induced TSS switching in T47D cells. Top and middle panel: the x-axis represents the distance to the AUG start codon, with the black bar denoting the RefSeq-annotated 5’UTR. The TSS cluster between 75-114 nt is expressed in both conditions, but has significantly different usage under hypoxia. Lower left panel: Bar plot of total transcript expression (all 5’UTR isoforms). Lower right panel: Bar plots summarizing 5’UTR isoform expression within each TSS cluster (nucleotide positions relative to the AUG start codon indicated below). The percentage of transcript expression for TSS cluster relative to the total transcript expression is indicated for each condition above bar plots, together with t-test p-values comparing isoform expression. Error bars indicate standard deviation. **f)** Same as **e**, but an example of a transcript with a qualitative hypoxia-induced TSS change in T47D cells. The TSS cluster between 27-37 nt is expressed only under normoxic conditions. **g-j)** An example of how change-point analysis was used to identify 5’UTR sequence regions in NM_000387 (SLC25A20) that are enriched, or depleted as a result of hypoxia-induced TSS switching seen in (**g**). Change-points define 5’UTR sequence windows with differential enrichment or depletion under hypoxia (**h**). The degree of isoform switching can be scored by taking the maximum difference in isoform expression across identified 5’UTR isoforms (TSS switch score) (**i**), demonstrating a large difference in isoform expression, even in the absence of changes in overall transcript levels (**j**). **k)** Barplot displaying categories of 5’UTR sequences enriched under hypoxia identified by change-point analysis. **l)** Venn diagram showing the overlap of transcripts with significant hypoxia-induced TSS switching identified in T47D and H9 cells (top panel). Of the transcripts with TSS switching in both cell lines (n = 586), comparison of change-point enrichments identified n = 219 transcripts with the same pattern of 5’UTR isoform switches in both cell types. Heatmap shows z-scores of the relative enrichment or depletion in each sequence segment defined by change-points for these 219 transcripts. Note that segments are not necessarily of equal lengths, but are visualized on a meta-transcript scale. **m)** Venn diagram showing the overlap of genes (note that a number of genes have several transcript isoforms) that undergo the same pattern of hypoxia-induced TSS switching in both T47D and H9 cells, and known transcriptional targets of HIF1α (from Schödel *et al*.^30^ and Sugimoto *et al*.^31^).

As TSS switching was often complex and involved multiple 5’UTR isoforms (**Fig 1g**), we employed change-point analysis^29^ to identify 5’UTR sequence segments (defined as the regions before, between, and after the change-points [short, middle, and long]) that are enriched or depleted under hypoxia, and used these to assign a score describing the extent of isoform switching (termed the TSS switch score, see methods; **Supplemental File S3**). For instance, for SLC25A20 (**Fig 1g**), change-points were identified 58 and 91 nucleotides (nt) upstream of the start codon (**Fig 1h**) and the relative expression of the resulting segments indicated a 20% increase of the shorter isoforms in hypoxia and concomitant depletion of isoforms longer than 91 nt (**Fig 1i**). The resulting TSS switch score of ∼49 (**Fig 1i**) indicates a high degree of 5’UTR remodeling in the absence of changes in corresponding mRNA levels (**Fig 1j**). We next applied this method to all transcripts with significant changes in TSS usage between hypoxia and normoxia in T47D or H9 cells, which revealed that TSS scores tended to be larger in T47D cells (**Fig S1g**). Furthermore, in T47D cells, hypoxia-enriched 5’UTR variants were more likely to be the shortest or longest, whereas in H9 cells, 5’UTRs were predominantly lengthened (**Fig 1k**). Moreover, hypoxia-enriched 5’UTR sequences were more GC-rich in T47D cells with increasing TSS switch scores (**Fig S1h**); and a similar pattern was observed in the H9 cells at lower (> 15) but not higher (> 50) TSS switch scores (**Fig S1i**), possibly reflecting the distinct pattern of gains of shorter vs. longer isoforms between the cell types (**Fig 1k**).

Out of all transcripts that switched TSS under hypoxia, 586 (∼11%) were shared across cell types. Among these, 219 (∼37%) exhibited highly similar changes in 5’UTR isoform expression (**Fig 1l**) and were significantly enriched for gene ontologies, including hypoxia response, glucose metabolism, cell cycle and proliferation, and chromatin remodelling (**Fig S1j**). This suggests that shared TSS switching events impact core hypoxia-related processes. Interestingly, of genes with similar switching in both cell types, fewer than 2% are known transcriptional targets of HIF1α^30,31^ (**Fig 1m**). Together, these findings reveal pervasive hypoxia-induced remodeling of 5’UTRs occurring in a largely HIF1-independent manner.

### TSS switching is associated with translational reprogramming under hypoxia

Reprogramming of mRNA translation is a critical component of cellular adaptations to low oxygen^2,5,18,32–34^. We next examined the impact of 48 (T47D cells) or 24 (H9 cells) hours of hypoxia (1% O_2_) on the translatome using polysome-profiling^35^. These conditions associated with stabilization of HIF1α, suppression of mTOR signaling, and induction of ISR, as evidenced by reductions in 4E-BP1 phosphorylation and increases in eIF2α phosphorylation, respectively (**Fig 2a-b**). Furthermore, as expected^2^, hypoxia suppressed global mRNA translation as illustrated by an 80-85% reduction in the polysome/monosome ratios in both cell types (**Fig 2c-d**).

**Figure 2:**
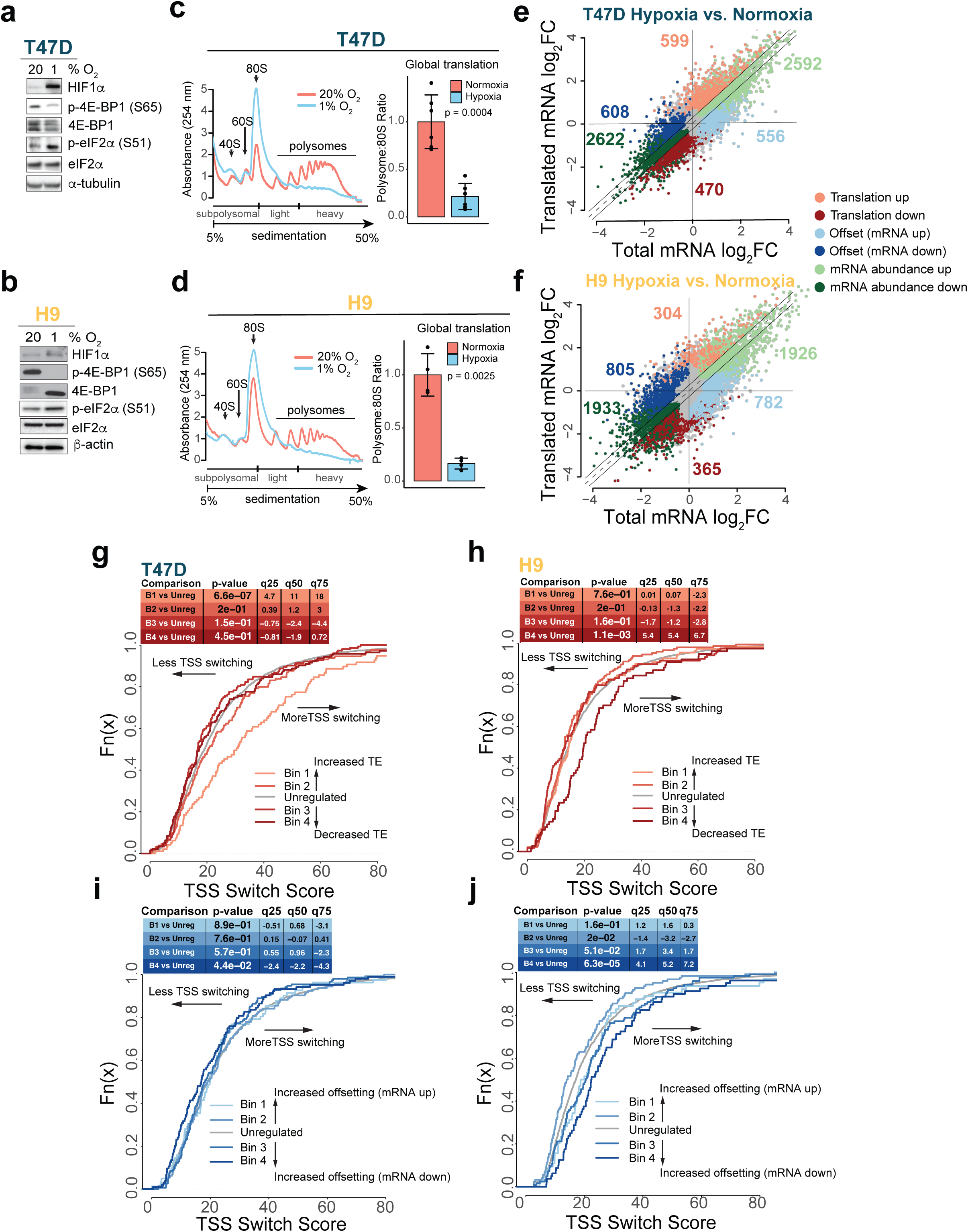
TSS switching is associated with translational reprogramming under hypoxia. **a-b)** Immunoblots of indicated proteins and cell types under normoxia and hypoxia. **c-d)** Polysome tracing and global translation change in hypoxia-treated T47D (**c**) and H9 (**d**) cells compared to normoxia. Polysome-associated mRNAs are considered those associated with >3 ribosomes. Global change in translation was quantified as the ratio between the area under the curve for polysomes and 80S monosomes in each condition (T47D: n = 6; H9: n = 4). Error bars indicate standard deviation. Significance determined by two-tailed t-test. **e-f)** Scatter plot of polysome-associated mRNA vs. total mRNA log_2_ fold changes in T47D (**e**) and H9 (**f**) cells (hypoxia vs. normoxia). Genes are colored according to their mode of regulation assigned by the anota2seq algorithm (FDR < 0.15). The number of regulated mRNAs in each category is indicated in corresponding colors. **g-h)** The 400 most up and downregulated genes were separated into four bins based on the quartiles of the fold changes in translation efficiency (determined by anota2seq analysis) for T47D (**g**) and H9 (**h**) cells. Empirical distribution functions (eCDFs) were used to compare the TSS switching scores (determined by change-point analysis) across the four bins. The set of background genes (i.e. not in bins) is also indicated (grey line). Significant shifts in TSS switch scores between each bin compared to background were assessed using Wilcoxon rank-sum tests. P-values and the magnitude of shifts at each quartile (q25-75) are indicated. Right-shifted curves indicate the sets of translationally regulated genes with more extensive TSS switching under hypoxia compared to unregulated genes. **i-j)** Same as **g-h** relating TSS switching scores to translational offsetting in T47D (**i**) and H9 (**j**) cells.

In addition to a global decrease in protein synthesis, transcript-selective changes in translation efficiency shape the newly synthesized proteome under hypoxia. To identify these hypoxia-induced changes in the translatome, we compared levels of total and heavy polysome-associated mRNA (> 3 ribosomes) quantified by RNA sequencing using the anota2seq algorithm^36^. In both cell lines non-congruent alterations in mRNA levels and polysome-association were observed for ∼2000 genes (**Fig 2e-f, Supplemental File S4**). This discordance arises due to changes in polysome-association of mRNAs not accompanied by corresponding alterations in their total level (“translation”; shown in red), or to changes in total mRNA levels that are translationally offset such that their polysome-association is unaltered (“offsetting”; shown in blue). Importantly, the “translation” mode is expected to result in altered protein levels, while “offsetting” opposes alterations in protein levels despite changes in mRNA abundance^37^. Notably, mRNAs encoding proteins implicated in the regulation of translation were induced but translationally offset under hypoxia in both T47D and H9 cells (**Fig S2a-b**). This included TOP mRNAs whose translation is mTOR-sensitive^23–25^ (**Fig S2c-d**).

To assess if 5’UTR remodelling associates with changes in translation, we compared TSS switch scores from nanoCAGE sequencing (**Fig S1g**) to changes in translation (**Fig 2e-f**). This revealed that high TSS switch scores associated with increased translation in T47D cells (**Fig 2g**) and suppressed translation in H9 cells (**Fig 2h**). While there was not a strong relationship between TSS switching and translational offsetting in T47D cells (**Fig 2i**), TSS switching associated with translational offsetting for mRNAs with decreased levels in H9 cells (**Fig 2j**). Together, these findings suggest that hypoxia-induced TSS switching impacts the translatome.

### Changes in the hypoxic translatome depend on multiple pathways and 5’UTR features

Translation is thought to largely be regulated by interactions between 5’UTR features and the translational machinery during the rate-limiting initiation step^21,38^. In yeast, TSS switching can lead to gain or loss of uORFs in 5’UTRs of a subset of mRNAs thus altering translation during meiosis or ER stress^39,40^. To better understand the contribution of TSS switching to translatome changes under hypoxia, we sought to first identify 5’UTR features and pathways contributing to hypoxia-associated translation (**Fig 3a** “Model 1”), and then assess the impact of TSS switching on the adaptive hypoxic translatome (**Fig 3a** “Model 2”). To achieve this, we developed a method that identifies variables explaining hypoxia-associated alterations in translation (see methods). The resulting networks represent regulatory nodes (i.e. 5’UTR features, pathways regulating mRNA translation, TSS switching, etc.) and edges indicating covariance between nodes (**Fig 3a-e**).

**Figure 3:**
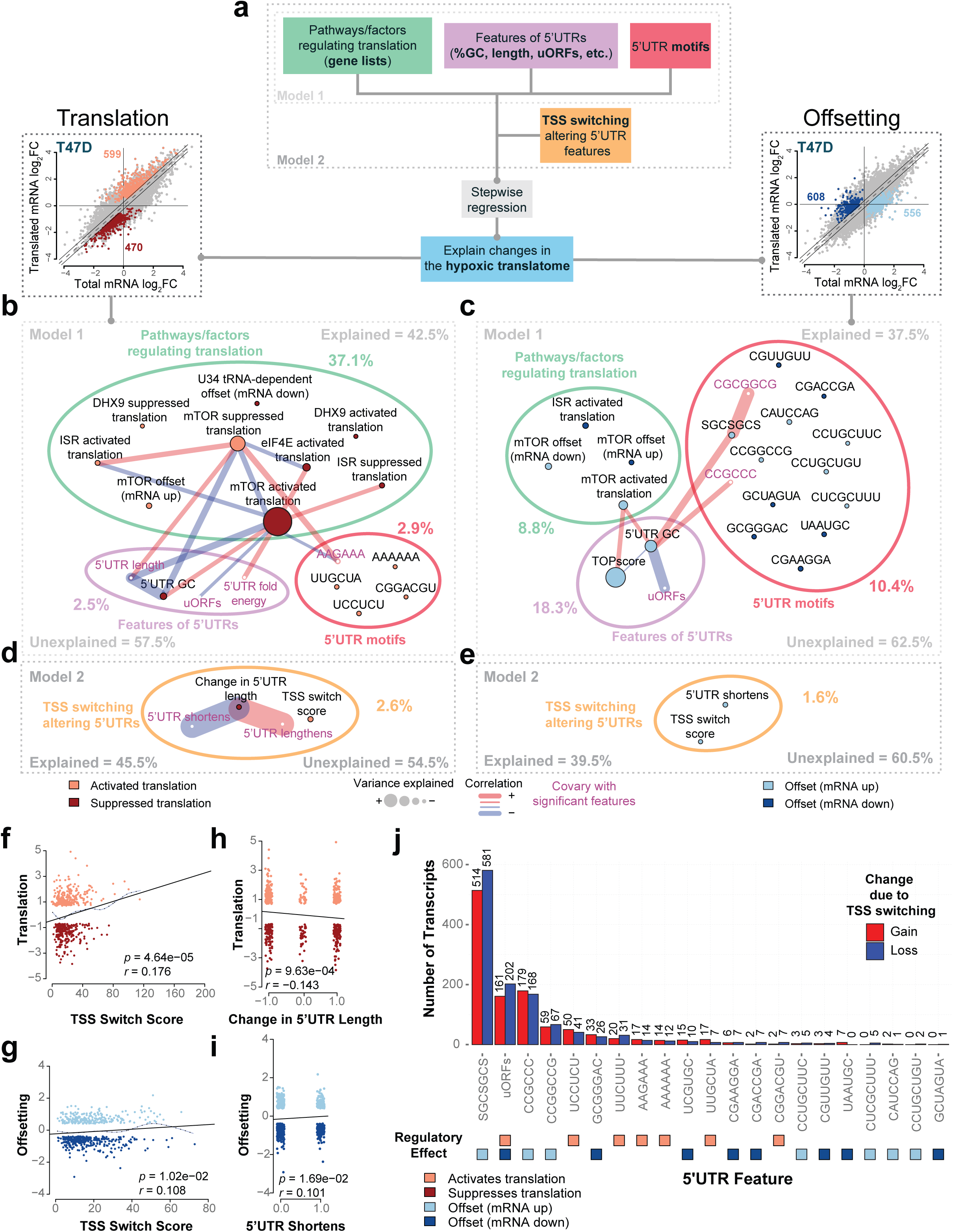
TSS switching alters regulatory 5’UTR features and shapes the hypoxia-induced translatome. **a)** Schematic describing translatome modeling. To identify features or pathways involved in hypoxic translatome remodelling, changes in translation efficiency or offsetting were modelled using known signatures of genes translationally regulated downstream of different pathways or factors (green), known mRNA features of 5’UTRs characterized by nanoCAGE sequencing in hypoxia-treated T47D cells (purple), and *de novo* 5’UTR motifs (pink) (Model 1). The impact of TSS switching was then assessed by adding signatures describing 5’UTR alterations to the modelling (Model 2). **b)** Network plot displaying the results of Model 1. The total percentage of changes in translation efficiency explained and unexplained are indicated. Also indicated are the percentages of translation changes explained by different input categories. Connections between features indicate substantial correlations. Colors of the nodes indicate if the feature is associated with translation activation or suppression under hypoxia. **c)** Same analysis as **b,** but modeling changes in translational offsetting induced by hypoxia in T47D cells. **d)** Selection of the full network plot for Model 2, displaying the additional impact of adding TSS-switching signatures to Model 1 in explaining changes in translation efficiency under hypoxia in T47D cells. **e)** Selection of the full network plot for Model 2, displaying the additional impact of adding TSS-switching signatures to Model 1 in explaining changes in translational offsetting under hypoxia in T47D cells. **f)** Scatterplot comparing the TSS switching score (indicative of altered 5’UTRs) vs. the change in translation efficiency that occurs under hypoxia. Pearson’s correlation coefficient and p-values indicated. **g)** Same as (**f**) for translational offsetting. **h)** Scatterplot comparing changes in 5’UTR length (resulting from TSS switching) vs. the change in translation efficiency that occurs under hypoxia. Pearson’s correlation coefficient and p-values indicated. **i)** Scatterplot comparing 5’UTR shortening events vs. the change in translational offsetting that occurs under hypoxia. Pearson’s correlation coefficient and p-values indicated. **j)** Barplot indicating the number of transcripts that gain or lose 5’UTR elements identified in either Model 1 or Model 2 in T47D cells. Transcripts were considered if more than 10% of the expressed pool of 5’UTR isoforms gained or lost the 5’UTR element. The translation mode associated with each element is indicated by colored squares below.

For genes with altered translation efficiency under hypoxia, modeling included previously established translational signatures (a total of 11 including mTOR, ISR, eIF4E2/4EHP, DAP5, etc.)^22,41–51^, 5’UTR features derived from the most abundant 5’UTR-isoform detected under hypoxia (including length, GC content, folding energy, the presence of uORFs, and known motifs), as well as *de novo* identified 5’UTR motifs (complete list of input variables is provided in **Supplemental File S4**) (**Fig 3a**). In T47D cells, the included variables explained 42.5% and 37.5% of hypoxia-induced changes in translation (**Fig 3b**) and offsetting (**Fig 3c**), respectively. As expected^5,18,20^, we found that mTOR suppression and ISR activation play prominent roles in hypoxia-associated alterations in translation, and are anti-correlated (i.e., genes translationally activated by ISR are in general suppressed by mTOR^46^, and vice versa) (**Fig 3b, Supplemental File S4**). Interestingly, hypoxia-induced changes in translational offsetting were also partially mTOR- and ISR-dependent (**Fig 3c, Supplemental File S4**). In addition, U34 tRNA modification-^49^ and RNA helicase DHX9-associated^41^ signatures explained changes in translation independently of ISR and mTOR suggesting a previously unappreciated role of these factors in translational regulation under hypoxia. Interestingly, the eIF4E-dependent signature also explained hypoxia-dependent translation independently of mTOR (**Fig 3b**). We also identified 5’UTR features associated with changes in the translatome under hypoxia (**Fig 3b-c, Supplemental File S4**). For instance, 5’UTR GC-content independently explained changes in translation and offsetting, with higher 5’UTR GC being associated with suppressed translation under hypoxia (**Fig 3b-c, S3a-b**). Furthermore, mRNAs harboring GC-rich 5’UTRs were translationally suppressed under hypoxia and overlapped with mTOR translational signatures (**Fig 3b**). In contrast, translation of mRNAs with longer, AU-rich, and/or uORF-containing 5’UTRs was induced upon mTOR inhibition and ISR induction under hypoxia, consistent with previous studies^23,46^ (**Fig 3b**). In agreement with the observed gene ontology enrichments (**Fig S2a-c**), the TOPscore^25^ (summarizing TOP motifs across 5’UTR isoforms) was the strongest predictor of offsetting (**Fig 3c**). Accordingly, this approach accurately captures known pathways and 5’UTR features mediating selective regulation of mRNA translation.

In addition to previously characterized regulatory features of 5’UTRs, we identified an AAGAAA motif associated with translational activation under hypoxia in T47D cells that was correlated with mTOR-sensitive translation, as well as a multitude of other sequence motifs in 5’UTRs associated with changes in translation and offsetting under hypoxia that did not co-vary (< 10%) with known translational signatures (**Fig 3b-c, Fig S3a-b**). For instance, the SGCSGCS (S = C or G) motif associated with translational offsetting. Some of these motifs are predicted to interact with RNA binding proteins (RBPs) and are largely distinct between translation and offsetting modes of regulation (**Supplemental File S4**). A number of these motifs were also enriched in mRNAs encoding factors involved in WNT signaling, cell adhesion, and angiogenesis (**Supplemental File S5**).

In H9 cells, 50.1% and 42.2% of the variance in hypoxia-associated translation and offsetting, respectively, were explained using the abovementioned analysis (**Fig S4a-b**). As in T47D cells, suppression of mTOR and activation of the ISR accounted for the greatest proportion of changes in translation efficiencies. In addition, signatures of eIF4E phosphorylation, eIF4GI, and DAP5, were also associated with hypoxia-induced changes in the translatome of H9 cells (**Fig S4a-b**). Interestingly, the 5’UTR motifs that were identified to modulate translation in T47D and H9 cells under hypoxia were largely distinct (**Fig 3b-c, Fig S4a-b, Fig S5a-b, Supplemental File S4**). This suggests that translational regulation in different cell types may be mediated via distinct repertoires of *trans*-acting factors including RBPs. Indeed, many RBPs with predicted binding to identified 5’UTR motifs were distinct between cell types, while others, like SRSF1, were shared (**Supplemental File S4**). A number of motifs identified in H9 cells were also significantly enriched among transcripts related to WNT signaling and cell migration similar to T47D, further suggesting that these processes may be translationally regulated under hypoxia (**Supplemental File S5**).

Overall, these findings confirm that suppression of mTOR and activation of the ISR play a pivotal role in translatome remodelling under hypoxia while implicating 5’UTRs features including length, GC content, folding energy, and the presence of uORFs. Furthermore, we identified several factors (e.g., U34 tRNA modification and DHX9) and 5’UTR motifs (e.g., the SGCSGCS motif) associated with changes in the hypoxic translatome that have not been previously reported, suggesting currently unknown mechanisms of translational control in hypoxia.

### TSS switching alters regulatory 5’UTR features and shapes the hypoxia-induced translatome

TSS switching alters 5’UTR features and associates with translational regulation. Hence, we next sought to determine whether TSS switching could independently explain changes in the hypoxic translatome (“Model 2”, **Fig. 3a,d-e**). In T47D cells, hypoxia-associated 5’UTR remodeling increased the explained proportion of variance to 45.5% and 39.5% for translation and offsetting modes of regulation, respectively (**Fig 3d-e**, full models shown in **Fig S3c-d**). Of note, alterations in 5’UTR length and the TSS switch score independently explained hypoxia-associated translation and offsetting modes of regulation (**Fig 3d-e**). Importantly, the independent contribution from TSS switching was comparable to that of ISR, and the combined independent effects of 5’UTR length, GC content, uORFs, and folding energy for translation (**Fig 3b-e, Supplemental File S4**). Higher TSS switch score (indicative of more extensive TSS switching) was associated with translational activation (**Fig 2g**, **Fig 3f**) and offsetting (**Fig 3g**). Furthermore, alterations in the length of 5’UTRs were also associated with changes in both translation and offsetting under hypoxia (**Fig 3h-i**). In H9 cells, including TSS switching explained 51.4% and 43.4% of hypoxia-induced alterations in translation and offsetting regulatory modes, respectively (**Fig S4c-d,** full models in **S5c-d**). Changes in TOP motifs (i.e., the difference in TOPscore^25^ between hypoxia and normoxia) explained 2.1% of the variance in translation (**Fig S4c and e, Fig S5c**), indicating that gain or loss of TOP motifs due to TSS switching altered the translatome under hypoxia in H9 cells (**Fig S4c and e**). Furthermore, 5’UTR lengthening associated with mRNAs whose levels were reduced but translationally offset under hypoxia (**Fig S4d and f, Fig S5d**).

Unlike specific 5’UTR features, such as TOP motifs or inhibitory uORFs, which impact translation efficiency with an expected directionality, TSS switching would be expected to both increase and decrease translation by causing the loss of a specific feature in some transcripts and the gain in others. A caveat of our approach is that it favours identification of features associated with either increased or decreased translation efficiency. As a consequence, we posit that the extent to which TSS switching regulates translation by modifying 5’UTR features is underestimated. Accordingly, we next assessed how the identified 5’UTR motifs (**Fig. 3b-c, S4a-b**) are impacted by TSS switching. In T47D cells, 514 transcripts gained one or more SGCSGCS motif (>10% change in the 5’UTR with a motif was used as a threshold for gain or loss), while 581 transcripts lost this motif (**Fig 3j**). Similarly, uORFs were gained or lost for 161 and 202 transcripts, respectively, leading to both activated and suppressed translation. Consistent with the finding that longer 5’UTR isoforms were most often enriched under hypoxia in H9 cells (**Fig 1k**), TSS switching more often resulted in the gain of 5’UTR regulatory elements (**Fig S4g**). For example, in H9 cells, 278 and 79 transcripts gained or lost uORFs, respectively, while the CCCUGC motif associated with translational suppression was gained and lost in 130 and 38 transcripts, respectively (**Fig S5g**).

Other identified motifs were also gained or lost in hundreds of transcripts, likely leading to both enhanced and suppressed translation and offsetting. Collectively, our analyses suggest that hypoxia-associated alterations in translational efficiency are driven by a myriad of 5’UTR features that can be altered due to TSS switching, in concert with remodeling of the translational machinery via e.g., suppression of mTOR and ISR induction. TSS switching is therefore a previously unappreciated mechanism that impacts the translatome under hypoxia.

### Hypoxia-induced TSS switching is associated with altered H3K4me3 and changes in nucleosome context

We next interrogated mechanisms driving TSS switching under hypoxia. TSS switching of HIF1α-target genes was previously described in VHL-null RCC4 renal cell carcinoma cells, where it was shown that these changes in TSS correlate with alterations in translation^31^. Although we also observed TSS switching for some transcriptional targets of HIF1α, a large proportion of the TSS switching events in both T47D and H9 cells occurred in the absence of changes in overall transcript abundance (**Fig 1d**). Furthermore, less than 2% of the equivalent TSS switching events between cell types were known HIF1α targets (**Fig 1m**), suggesting that hypoxia-induced TSS switching may not be directly linked to HIF1. Hypoxia can broadly regulate the epigenome, via alterations in histone methylation due to the loss of activity of demethylases including KDM5A^6,8^. Because H3K4me3 marks TSSs, and KDM5 activity is reduced in hypoxia, we examined whether alterations in H3K4me3 correlate with TSS switching. As hypoxia leads to increased H3K4me3 (**Fig 4a**), we assessed differences in H3K4me3 distribution around TSSs using H3K4me3-ChIPseq (**Fig S6a-b**). This revealed hypoxia-associated differences in H3K4me3 distributions around TSSs for almost all protein-coding genes detected by nanoCAGE sequencing (94% and 98% at FDR < 0.01 in T47D and H9, respectively) (**Fig 4b, Supplemental File S6**), that fell into three categories: downstream or upstream shifts, and other alterations without a dominant directionality (e.g., genes in **Fig 4c** that also switch TSS in T47D cells upon hypoxia; **Supplemental File S1**). The proportions of these changes in H3K4me3 differed dramatically between cell lines (**Fig 4d**) and partly mirrored observed hypoxia-induced alterations in 5’UTR sequence segments identified by change-point analysis (**Fig 1k).** Although significant TSS switching events were associated with all three categories of H3K4me3 changes in both cell lines (**Supplemental File S1**), these findings are consistent with the deposition of H3K4me3 under hypoxia being dependent on the existing epigenetic landscapes and therefore diverging between cell types.

**Figure 4:**
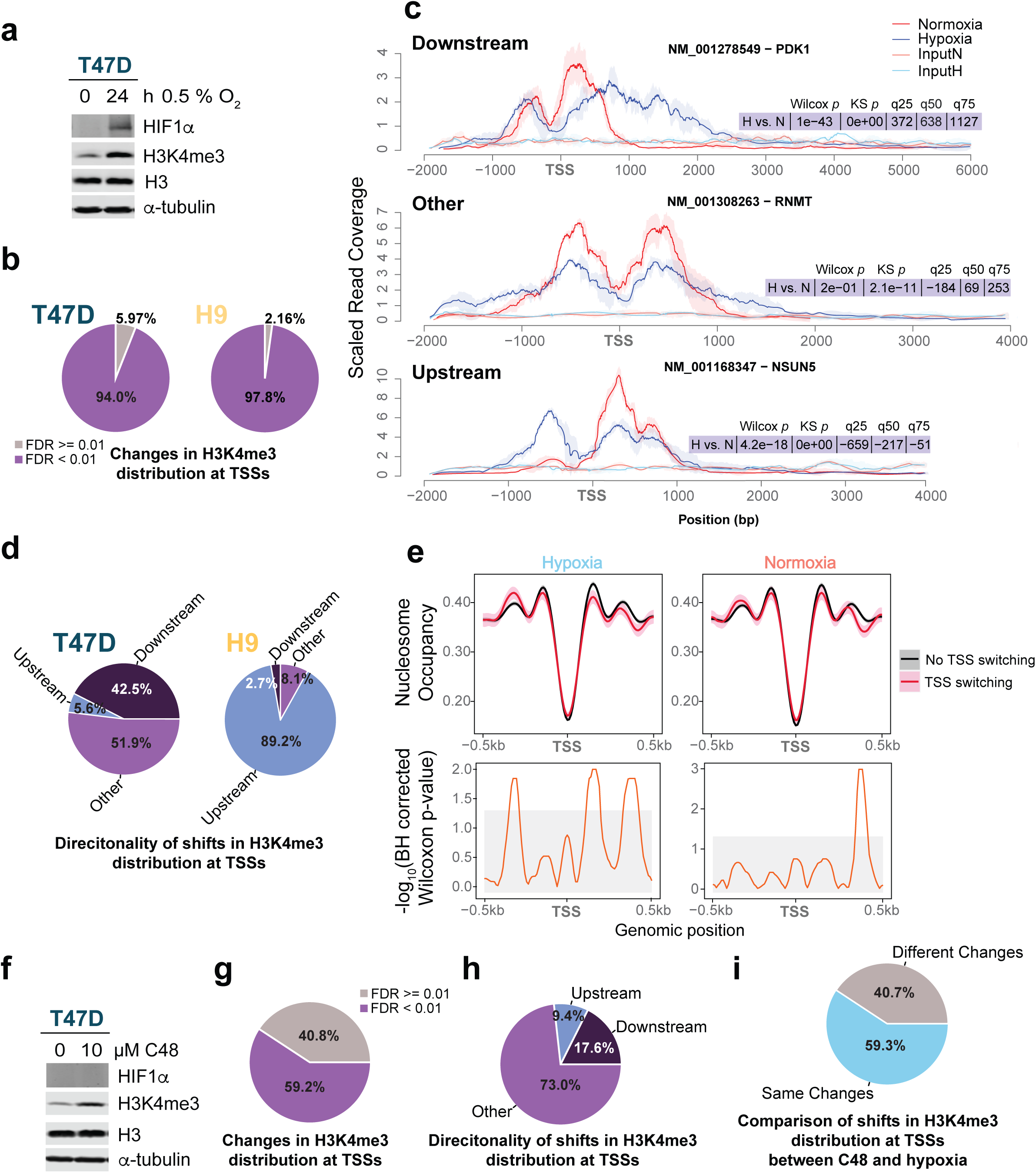
Hypoxia-induced TSS switching is associated with altered H3K4me3 and changes in nucleosome context. **a)** Immunoblots of HIF1α and H3K4me3 following 0 and 24 hrs of hypoxia in T47D cells. H3 and -tubulin were used as loading controls. **b)** Changes in the distribution of the H3K4me3 marks at TSSs were detected using a Kolmorogov-Smirnov test. The proportion of TSSs for protein-coding genes with significant changes beyond a threshold of FDR < 0.01 are shown. N = 3 for both T47D and H9 cells, and the mean of the replicates was used in all comparisons. **c)** Examples of genes with downstream, upstream, or other (in this case, a broadening) change in the distribution of H3K4me3 around the TSS that is accompanied by significant TSS switching under hypoxia in T47D cells. The Wilcoxon and Kolmogorov-Smirnov p-values comparing the distributions are indicated, as well as the directionality and magnitude of the shifts in the distribution at the quartiles. **d)** Summary of directional shifts in H3K4me3 distributions detected by two-tailed Wilcoxon signed-rank test. Directionality was assigned by comparing shifts in the distribution between hypoxia and normoxia at the quartiles of the H3K4me3 distribution for each locus. The proportion of TSSs for protein-coding genes with changes (FDR < 0.01) is shown. N = 3 for both T47D and H9 cells, and the mean of the replicates was used in all comparisons. **e)** Top panels show nucleosome occupancy determined using NucleoATAC^52^ analysis of ATAC-seq performed on T47D cells under hypoxia or normoxia (0.5% or 20% O_2_) for 48 h. Occupancy signal is anchored around the nanoCAGE-determined TSS under normoxia (most abundant TSS peaks). Lines indicate the mean and shaded areas the bootstrap confidence interval (1000 iterations, CI = 0.95). Bottom panels indicate the adjusted Wilcoxon rank-sum p-values between nucleosome occupancy signals across bins. Grey-shaded areas represent an FDR threshold of 0.05. **f)** Immunoblots of HIF1α and H3K4me3 following 24 hrs of 10 μM C48 or DMSO treatment in T47D cells. H3 and -tubulin were used as loading controls. **g)** Changes in the distribution of the H3K4me3 marks at TSSs in C48-treated T47D cells, as in (**b**). N = 4, and the mean of replicates was used in comparisons. **h)** Significant directional shifts in H3K4me3 distributions in C48-treated T47D cells, as in (**d**). N = 4 and the mean of the replicates was used in comparisons. **i**) Comparison of the directionality of significant changes in H3K4me3 distribution around TSSs between T47D cells treated with hypoxia vs. normoxia, and C48 vs. DMSO. Changes were considered equivalent if the directionality of shifts in the distribution at each quartile were the same between treatment conditions.

To examine if TSS switching associates with changes in nucleosome context around TSSs we performed ATAC-seq in T47D cells under normoxia and hypoxia (**Fig S6c**), followed by analysis of nucleosome positioning and occupancy using the NucleoATAC algorithm^52^. This indicated that genes undergoing TSS switching under hypoxia had a decrease and increase in occupancy at the +1 and -2 nucleosomes, respectively, relative to normoxia (**Fig 4e**). Although TSS switching events were associated with changes in H3K4me3 distribution and distinct nucleosome remodelling, the hypoxic response is complex and impacts many cellular processes, including additional epigenetic alterations^6–8^. To focus on the effects of H3K4me3 we treated T47D cells with the selective KDM5 inhibitor C48 (10 µM; 24 h), which led to a similar increase in H3K4me3 as 24 hrs of 0.5% O_2_ while not being accompanied by HIF1α stabilization (**Fig 4a, f**). Using ChIP-seq (**Fig. S6d)** we observed that C48 treatment resulted in significant changes in H3K4me3 distribution around the TSS of 52.9% of detected protein-coding genes (**Fig 4g**) categorized into downstream shifts (17.6%), upstream shifts (9.4%), and other alterations (73.0%) (**Fig 4h**). Comparing C48 treatment to hypoxia, we found that more than 59% of H3K4me3 alterations had the same directionality (downstream, upstream, or other) (**Fig 4i**), confirming that KDM5 inhibition at least partially reproduced a large proportion of hypoxia-induced H3K4me3 remodelling.

### Inhibition of KDM5 induces TSS switching that remodels 5’UTRs and is associated with proteome changes

To further establish the role of H3K4me3 in mediating hypoxia-induced TSS switching, we performed nanoCAGE sequencing (**Fig S7a**) in C48 or vehicle (DMSO)-treated T47D cells (**Fig 5a-b**), which identified TSS switching for > 3,000 transcripts (**Fig 5c-d**, **Supplemental File S1**). TSS switch scores were lower in C48 treated vs. hypoxia-exposed cells, suggesting that mechanisms other than H3K4me3 may tune the magnitude of the isoform switching under hypoxia (**Fig S7b**). A comparison of 5’UTR sequence segment-enrichments from change-point analysis showed that 682 transcripts had the same enrichments with C48 treatment as under hypoxia (28% of hypoxia-associated TSS switching) (**Supplemental File S3**). A Monte Carlo simulation showed that this was a greater proportion than expected by chance (**Fig 5e**). Of the 682 transcripts with the same TSS-switching, 32% and 79% did not exhibit changes in mRNA level under hypoxia and C48 treatment respectively (**Fig 5f**). Therefore, modulation of H3K4me3 caused by KDM5 inhibition is sufficient to alter TSS selection independent of hypoxic transcriptional programs.

**Figure 5:**
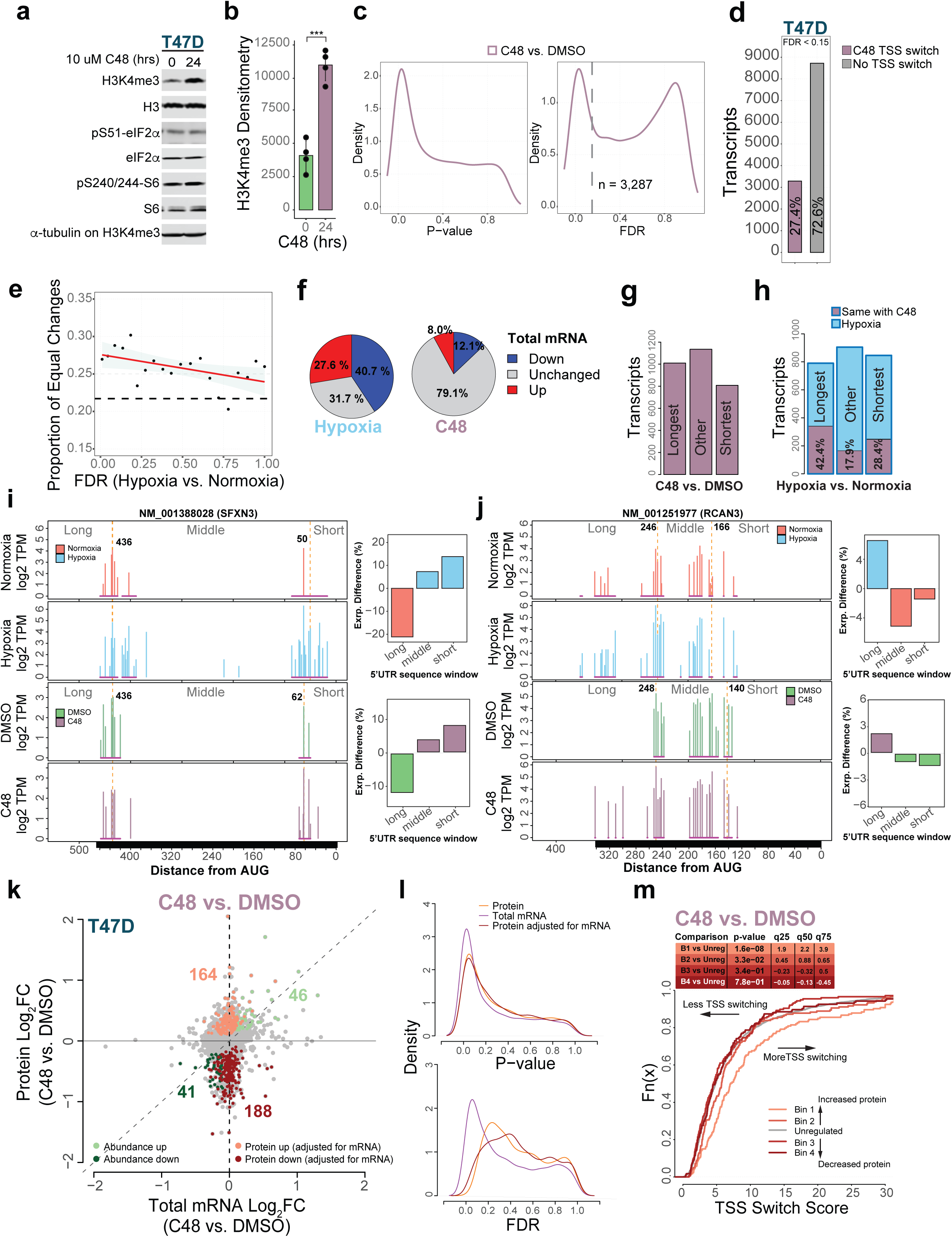
Inhibition of KDM5 induces TSS switching that remodels 5’UTRs and is associated with proteome changes. **a)** Immunoblots of H3K4me3, phosphorylated eIF2α (S51), and S6 (S240/244), from T47D cells treated with 10 μM C48 for 24 hrs, or DMSO (0 hr). H3, eIF2α, S6, and α-tubulin were used as loading controls. Representative blot shown (n = 4). **b)** Densitometry of immunoblots of H3K4me3 normalized to α-tubulin and H3 loading controls. Significance determined by two-tailed t-test (n = 4). **c)** Kernel density estimation P-value (top panel) and FDR (bottom panel) distributions for differential TSS usage between C48 and DMSO-treated T47D cells. Dotted grey line indicates an FDR threshold of 0.15. **d)** Barplot of protein-coding transcripts with significantly altered TSS usage between C48 and DMSO treatments (FDR < 0.15) in T47D cells. **e)** More significant TSS switching events under hypoxia are more likely to be recapitulated by C48 treatment. The FDR range for TSS switching under hypoxia was divided into ventiles and the proportion of equivalent changes in 5’UTR isoforms between hypoxia vs. normoxia and C48 vs. DMSO comparisons was determined. The dashed horizontal line marks the proportion of changes expected to be the same by chance (21.8%, estimated by Monte Carlo simulation). **f)** By comparing the 5’UTR sequence windows from change-point analysis of transcripts undergoing TSS switching under hypoxia vs C48 treatment, 682 transcripts with the same changes in 5’UTR isoform expression were identified. Pie charts indicate the proportions of these transcripts that have significantly (FDR < 0.15) increased (up), decreased (down), or unchanged overall expression between hypoxia and normoxia in T47D (left) and between C48 and DMSO (right). **g)** Barplot of categories of 5’UTR sequences enriched in transcripts with significant TSS switching after C48 treatment, identified by change-point analysis (n = 3,287). **h)** Barplot of categories of 5’UTR sequences enriched for transcripts with significant TSS switching between hypoxia and normoxia, identified by change-point analysis. The percentage recapitulated with C48 treatment is indicated. In total 30.1% of change-point-defined 5’UTR categorical changes under hypoxia were recapitulated by C48 treatment. **i)** 5’UTR isoform expression for NM_001388028 (SFXN3) in hypoxia and normoxia-treated (top), and C48 and DMSO-treated T47D cells (bottom). 5’UTR isoform changes within windows defined by change-point analysis are shown (right panels). Both hypoxia and C48 treatment enrich shorter 5’UTR isoforms. **j)** Same as (**i**) but for NM_001251977 (RCAN3), where both hypoxia and C48 treatment enrich longer 5’UTR isoforms. **k)** Scatter plot of protein (from GPF-DIA proteomics analysis) vs. total mRNA log_2_ fold changes in T47D cells (C48 vs. DMSO). Genes are colored according to their mode of regulation assigned by the anota2seq algorithm (FDR < 0.15), where protein up and down categories represent changes in protein level occurring independently of changes in mRNA level. Abundance up and down categories represent genes where changes in protein and mRNA levels are congruent. The number of regulated genes in each category is indicated in corresponding colors. **l)** Kernel density estimation P-value (top panel) and FDR (bottom panel) distributions for anota2seq analysis of changes in protein levels, total mRNA levels, and protein adjusted for mRNA levels between C48 and DMSO-treated T47D cells (n = 4). **m)** The 400 most up and downregulated genes were separated into four bins based on the quartiles of the fold changes in protein levels adjusted for mRNA levels (determined by anota2seq analysis) in T47D cells treated with C48. Empirical distribution functions (eCDFs) were used to compare the TSS switching scores (determined by change-point analysis) across the four bins. The set of all unregulated genes was used as the background (grey line). Differences in TSS score between each bin compared to background were assessed using Wilcoxon rank-sum tests. P-values and the magnitude of shifts at each quartile (q25-75) are indicated. Right-shifted curves indicate the sets of genes with more extensive TSS switching after C48 treatment compared to unregulated genes.

The requirement for all sequence segment-enrichments to be equivalent between hypoxia and C48-induced TSS switching is stringent and does not account for partial overlaps. We therefore also examined categorical changes. Enrichments of 5’UTR isoforms following C48 treatment (**Fig 5g**) resembled those triggered by hypoxia-induced TSS switching (**Fig 1k**). Of the transcripts that underwent significant TSS switching enriching for the longest and shortest 5’UTR isoforms under hypoxia, C48 treatment was found to result in the same pattern of enrichment for 42.4% and 28.4% of these mRNAs, respectively (**Fig 5h**). Overall, 30.1% of hypoxia-induced TSS switching was either fully or partially observed with C48 treatment. SFXN3 illustrates the similarity between hypoxia and C48-induced TSS switching as change-point analysis identified a relative loss of longer (> 436 nt) and increase of shorter 5’UTR isoforms (< 436 nt) under hypoxia (**Fig 5i**, top panels), which was mirrored by C48 treatment (**Fig 5i**, bottom panels). Similarly, under hypoxia, TSS switching increased levels of longer 5’UTR isoforms (> 246 nt) for RCAN3, and the same pattern of 5’UTR isoform expression was observed after C48 treatment (**Fig 5j**). Accordingly, inhibition of KDM5 induces extensive TSS switching that recapitulates a significant proportion of TSS alterations observed under hypoxia.

We next interrogated whether this altered pool of 5’UTR isoforms impacts the proteome under C48 treatment. Importantly, unlike hypoxia, C48 treatment did not alter mTOR or ISR signaling, as evidenced by the lack of changes in phosphorylation of S6 and eIF2α (**Fig 5a)**. We performed GPF-DIA proteomics in parallel with RNA-seq on the same C48 and DMSO-treated cells used to profile TSSs (**Fig S7d-e**). Using anota2seq^36^, we identified changes in protein levels not paralleled by corresponding changes in mRNA level following C48 treatment (n = 352, FDR < 0.15) (**Fig 5k-l**). These proteins were enriched for gene ontologies related to extracellular vesicles, protein modifications, and response to hypoxia, among others (**Fig S7f**). Similarly to hypoxia in T47D cells (**Fig 2g**), higher TSS switch scores following C48 treatment were associated with increased protein levels, independent of mRNA level (**Fig 5m**). Together, these findings demonstrate that KDM5 inhibition alone is sufficient to induce TSS switching that recapitulates a subset of hypoxia-induced switching and contributes to proteome alterations. Therefore, TSS switching-dependent remodeling of the 5’UTRome affects the proteome independently of changes in mRNA levels, and in the absence of alterations in mTOR or ISR signaling.

### Inhibiting H3K4me3 accumulation under hypoxia blocks TSS switching and decreases cellular fitness

We next tested the effects of attenuating H3K4me3 on hypoxia-induced TSS switching. To this end, we pre-treated T47D cells with DMSO or OICR-9429 (25 μM, 48 hrs), an inhibitor of the interaction between WDR5 and MLL-associated COMPASS H3K4 methyltransferases^53^, followed by exposure to hypoxia (0.5% O_2_) for an additional 48 h. OICR-9429 reduced H3K4me3 accumulation under hypoxia (**Fig 6a-b**). NanoCAGE mapping of TSSs (**Fig S7g**) revealed that addition of OICR-9429 under hypoxia leads to TSS switching for > 5,000 transcripts (FDR < 0.15) compared to hypoxia alone. 618 of the OICR-9429-induced TSS switching events showed sequence segment enrichments opposite to those observed under hypoxia, which was more than expected by chance (**Fig 6c-e**). In contrast to C48 treatment, treatment with OICR-9429 under hypoxia predominantly resulted in enriched expression of shorter 5’UTR isoforms (**Fig 6f**) and reversed hypoxia-associated enrichment for 57.6% and 26.0% of the longest and shortest transcripts, respectively (**Fig 6g**). For example, under hypoxia, TSS usage resulting in expression of shorter 5’UTR isoforms of TOP3A was diminished. However, upon addition of OICR-9429, expression of these shorter isoforms was restored (**Fig 6h**). Comparing the subsets of hypoxia-induced TSS switching events that were recapitulated with inhibition of KDM5 and blocked by OICR-9429 under hypoxia, we found a group of transcripts for which TSS selection appears to be highly dependent on H3K4me3 (n = 268) (**Fig 6i**). This subset was enriched for gene ontologies such as protein modifications, cell adhesion, and cellular ion homeostasis (**Fig S7h**), and included the nuclear import factor TNPO3, where both hypoxia and C48 treatment resulted in a relative depletion of longer 5’UTR isoforms, which was blocked by addition of OICR-9429 (**Fig 6j**).

**Figure 6:**
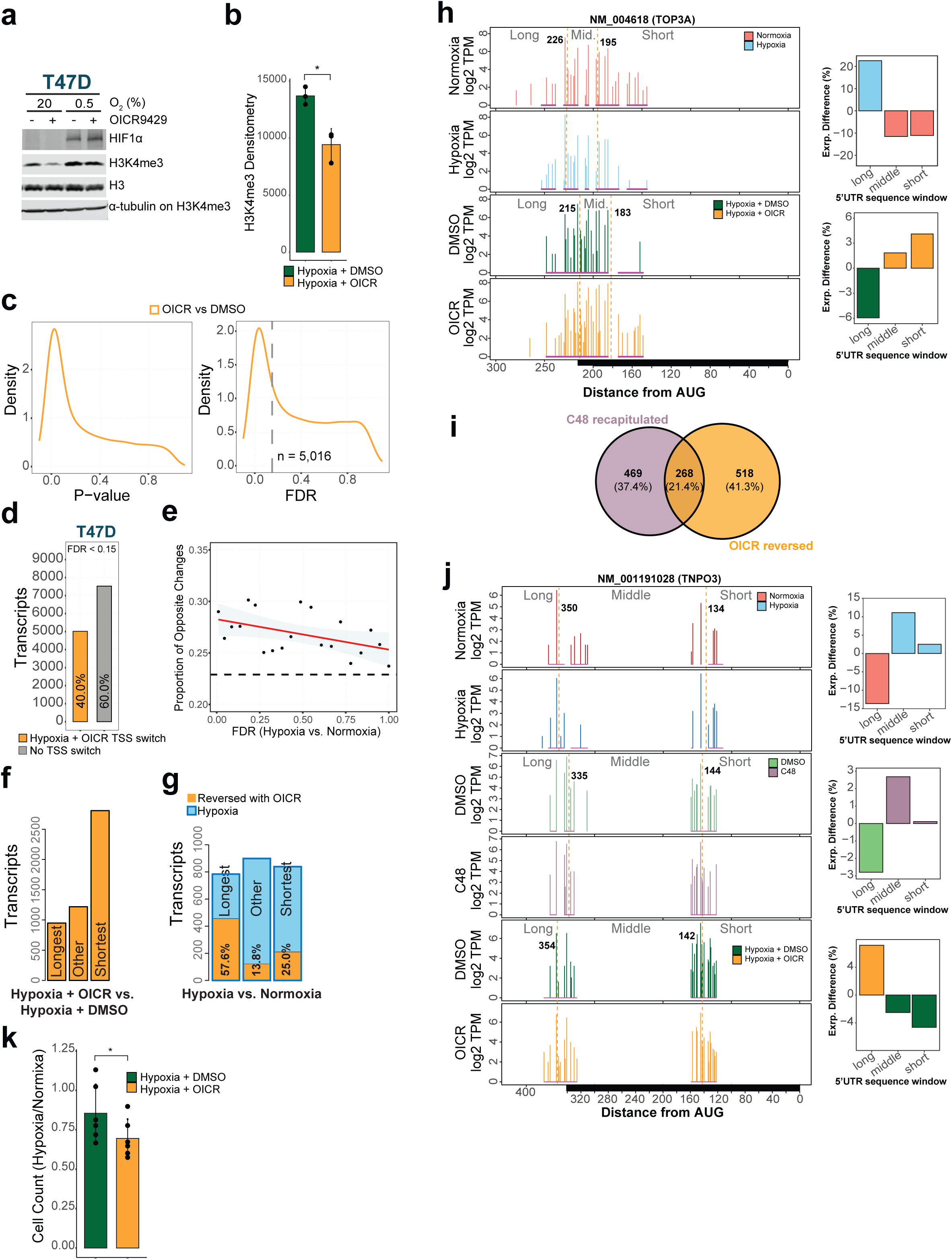
Inhibiting H3K4me3 accumulation under hypoxia blocks TSS switching and decreases cellular fitness. **a)** Immunoblots of H3K4me3 from T47D cells pretreated with 25µM OICR-9429 or DMSO for 48h and treated with hypoxia or normoxia, with the addition of 25µM OICR-9429 or DMSO for an additional 48h. H3 and α-tubulin were used as loading controls. HIF1α was used as a positive control for hypoxia. Representative blot shown (n = 3). **b)** Densitometry of immunoblots of H3K4me3 normalized to α-tubulin and H3 loading controls. Bars represent the mean and error bars indicate the standard deviation. Significance determined by two-tailed t-test (n = 3). **c)** Kernel density estimation P-value (left panel) and FDR (right panel) distributions for differential TSS usage in T47D cells between co-treatments of hypoxia and OICR-9429 or DMSO. Dotted grey line indicates an FDR threshold of 0.15). **d)** Barplot indicating the proportions of protein-coding transcripts with significantly altered TSS usage between T47D cells co-treated with hypoxia and OICR-9429 vs DMSO (FDR < 0.15) (n = 5,016). **e)** More significant TSS switching events under hypoxia are more likely to be reversed by OICR-9429 treatment. The FDR range for TSS switching under hypoxia was divided into ventiles and the proportion of opposite changes in 5’UTR isoforms between hypoxia + OICR-9429 vs. hypoxia + DMSO comparisons was quantified. The dashed horizontal line marks the proportion of changes that could be expected to be the same by chance (22.9%, estimated by Monte Carlo simulation). **f)** Barplot of categories of 5’UTR sequences enriched in transcripts with significant TSS switching after co-treatment with hypoxia and OICR-9429, identified by change-point analysis (n = 5,016). **g)** Barplot of categories of 5’UTR sequences from transcripts with significant TSS switching between hypoxia and normoxia, identified by change-point analysis. The percentage that was reversed with the addition of OICR-9429 is indicated. In total, 786 (31.1%) of change-point-defined 5’UTR categorical changes under hypoxia were reversed by OICR-9429. **h)** Quantification of 5’UTR isoforms of NM_004618 (TOP3A) in hypoxia and normoxia-treated (top), and hypoxia + DMSO or OICR-9429-treated T47D cells (bottom). Enriched and depleted 5’UTR sequence windows in both treatment comparisons are shown (right panels). The addition of OICR-9429 restores expression of shorter 5’UTR isoforms that were lost under hypoxia. **i)** Venn diagram showing the overlap of transcripts where hypoxia-induced TSS switching was recapitulated by C48 treatment, and reversed by OICR-9429 treatment in T47D cells. **j)** Quantification of 5’UTR isoforms of NM_001191028 (TNPO3) in hypoxia and normoxia-treated (top panel), C48 and DMSO-treated (middle panel), and hypoxia + DMSO or OICR-9429-treated T47D cells (bottom panel). Enriched and depleted 5’UTR sequence windows in all three treatment comparisons are shown (right panels). C48 treatment recapitulates the enrichment in expression of shorter 5’UTR isoforms that occurs under hypoxia while OICR-9429 reverses this effect. **k)** Trypan blue exclusion assays to quantify viable cells under hypoxia. T47D cells treated as in (**a**) were counted at endpoint. Bars represent the mean of the ratio of cell count in hypoxia vs. normoxia in OICR-9429 or DMSO-treated cells and error bars indicate the standard deviation. Significance determined by two-tailed t-test (n = 6).

Finally, to evaluate the importance of epigenetically-mediated TSS switching in cellular fitness under hypoxia, we measured proliferation of T47D cells under hypoxia, co-treated with a vehicle (DMSO) or OICR-9429. Addition of OICR-9429, which blocked close to 30% of hypoxia-induced TSS switching, resulted in a significant decrease in cell proliferation under hypoxia, relative to the same treatment under normoxia (**Fig 6k**). Taken together, these findings indicate that alterations to H3K4me3 under hypoxia account for a substantial proportion of TSS switching events, identifying a previously unknown role for this epigenetic modification in 5’UTR-determining TSS selection that contributes to cellular adaptations under hypoxia.

### TSS switching orchestrates adaptation to hypoxia by regulating availability of differentially translated mRNA isoforms

We next examined whether specific biological processes are regulated by TSS-switching. Gene ontology analysis of transcripts with similar hypoxia-induced TSS switching events between T47D and H9 cells (**Fig S1j**), and those with significant TSS switching in the individual cell lines revealed enrichment of metabolism-related terms (**Fig S8a**). To survive hypoxia, cells must switch from oxidative phosphorylation to glycolysis^54^ and, strikingly, many enzymes involved in glycolysis underwent significant TSS switching in hypoxic T47D cells (**Fig S8b**) including PDK1, which showed one of the most significant TSS switching events in both cell lines (**Fig 7a, Fig S8c, Supplemental File S1**). Under hypoxia, PDK1 phosphorylates Pyruvate Dehydrogenase (PDH), thus preventing entry of pyruvate into the citric acid cycle. This conserves pyruvate for other metabolic processes required for adaptation to hypoxia, including its reduction to lactate to regenerate NAD+ for subsequent rounds of glycolysis^54^. PDK1 transcription is induced under hypoxia by HIF1α. In addition, PDK1 is activated by ATP, NADH and CoA and inactivated by ADP, NAD+, pyruvate and CoA-SH^55^. As expected, hypoxia increased PDK1 protein levels and PDH1 phosphorylation at S232 (**Fig 7b**). Change-point analysis of PDK1 5’UTR isoform expression in T47D cells showed a loss of longer (> 87 nt) 5’UTRs and enrichment of shorter 5’UTR isoforms (**Fig 7c**). We confirmed expression of the short, hypoxia-inducible isoforms using 5’RACE (**Fig 7a**), and isoform-selective RT-qPCR (**Fig 7d**). In H9 cells, which are more glycolytic^2,56^, more PDK1 5’UTR isoforms were detected under normoxia than in T47D cells (**Fig S8c)**. Nevertheless, change-points and 5’UTR sequence segment enrichments were highly similar (**Fig S8d**), suggesting that hypoxia-induced TSS switching may be an important regulator of PDK1 expression.

**Figure 7:**
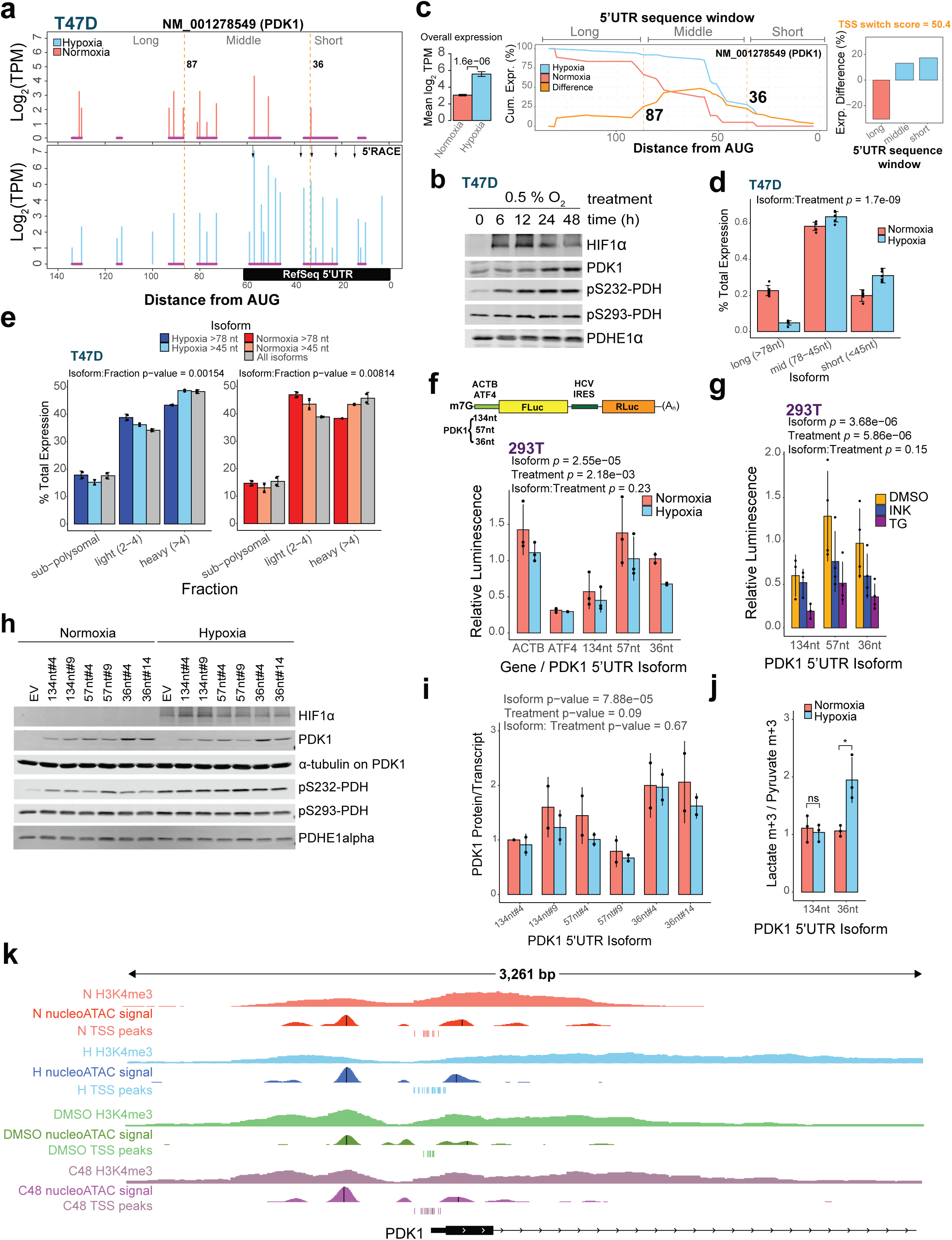
TSS switching orchestrates adaptation to hypoxia by regulating availability of differentially translated mRNA isoforms. **a)** Quantification of 5’UTR isoforms for NM_001278549 (PDK1) in hypoxia and normoxia-treated T47D cells. Same outline as **Fig 1e-f** but also indicating change-point-identified sequence windows by vertical orange lines and isoforms detected by 5’RACE in hypoxia-treated T47D by black arrows in the hypoxia condition. **b)** Immunoblot showing PDK1, PDHE1 and phosphorylated PDH (S323 and 293) with increasing time under hypoxia treatment in T47D cells. HIF1 is used as a positive control for hypoxia. **c)** Change-point analysis of NM_001278549 identifies change-points at positions 36 and 87 nucleotides upstream of the start codon (middle panel), which define shorter sequence windows with isoforms that are enriched under hypoxia and a longer window that is depleted (right panel). Difference in total transcript expression is also shown (left panel). **d)** Quantification of PDK1 5’UTR isoform expression relative to total transcript in hypoxia and normoxia-treated T47D cells using RT-qPCR. P-value shown is for the isoform:treatment interaction from a linear model with the design: % Isoform Expression ∼ Replicate + Isoform + Treatment + Isoform:Treatment, evaluating if the pattern of isoform expression differs between conditions. Bars represent the mean, and error bars indicate standard deviation (n = 4). **e)** Polysome occupancy of PDK1 5’UTR isoforms under hypoxia (left panel) and normoxia (right panel). RNA was isolated from each sucrose fraction separated by polysome fractionation and subjected to RT-qPCR. The proportion of PDK1 5’UTR isoforms measured by RT-qPCR in sub-polysomal, light (2-4), and heavy (>4) polysome fractions is shown, where the sum of all fractions for each mRNA is set to 100. P-value shown is for the isoform:treatment interaction from a linear model with the design: % Expression ∼ Isoform + Fraction + Isoform:Fraction, evaluating if the pattern of isoform abundance differs between fractions. Bars represent the mean and error bars indicate standard deviation (n = 2). **f)** Top panel: Schematic of m7G-capped bicistronic reporter mRNA harbouring the 5’UTR of PDK1 (134, 57 or 36 nucleotides), or the 5’UTR of beta-actin or ATF4, upstream of the firefly luciferase (FLuc) ORF and HCV IRES upstream of Renilla luciferase (Rluc) ORF. Bottom panel: Firefly relative to Renilla luciferase signal in normoxia- and hypoxia-treated 293T cells transfected with the reporter mRNA. P-values are provided for differences between translation of PDK1 5’UTR isoforms, differences translation depending on treatment, and the interaction between isoform:treatment (testing if isoforms are differentially translated between treatments) from a linear model with the design log2(Luminescence) ∼ Replicate + Isoform + Treatment + Isoform:Treatment. Bars represent the mean and error bars indicate standard deviation (n = 3). **g)** Firefly luciferase values relative to Renilla luciferase in DMSO, Thapsigargin (TG, 400 nM) and INK128 (INK, 50 nM) treated 293T transfected with the reporter mRNA containing PDK1 5’UTR isoforms. P-values are provided for the linear model with the same design as **f**. Bars represent the mean, and error bars indicate standard deviation. **h)** Immunoblot showing PDK1 and phosphorylated PDH (S323 and 293) under hypoxia treatment (24 hrs) in T47D cells with KO of endogenous PDK1 and re-expression of individual PDK1 5’UTR-isoforms. Two independent clones are shown per isoform. PDHE1 is used as a loading control and HIF1 is used as a positive control for hypoxia. **i)** The ratio of PDK1 protein and transcript levels in T47D cells with KO of endogenous PDK1 and re-expression of individual 5’UTRs after 24 hrs of hypoxia treatment. P-values are provided for differences between 5’UTR isoforms, differences depending on treatment, and the interaction between isoform:treatment from a linear model with the design Protein ∼ transcript + Isoform + Treatment + Replicate + Isoform:Treatment. Bars represent the mean and error bars indicate standard deviation (n = 2). **j)** Pyruvate conversion to lactate is enhanced upon expression of shorter 5’UTR isoforms in hypoxia. The ratio of labelled lactate m+3 / pyruvate m+3 from T47D cells with KO of endogenous PDK1 and re-expression of individual PDK1 5’UTR isoforms grown in hypoxia or normoxia for 24 hr. Metabolites were determined using stable isotope tracing by GC-MS. Bars represent the mean, and error bars the standard deviation (n = 3). Significant differences determined by two-tailed t-test . * indicates p-value < 0.05. **k)** H3K4me3, nucleosome occupancy, and TSS peaks for NM_001278549 isoform of PDK1 under hypoxia and normoxia, and with DMSO and C48 treatments. H3K4me3 was measured by ChIPseq, Reproducible TSS peaks measured by nanoCAGE are indicated. Nucleosome occupancy is displayed as the smoothed nucleoATAC signal. Vertical black bars indicate the dyad positions of nucleosomes determined using nucleoATAC.

We next examined whether translation of PDK1 5’UTR isoforms depends on oxygen availability by monitoring their distribution across a sucrose gradient under normoxia or hypoxia (**Fig 7e**, **Fig S8e**). This revealed that under hypoxia, the shorter (less than 45 nt) isoforms induced by TSS switching were enriched in heavier polysomes and therefore more efficiently translated relative to the longer (more than 78 nt) 5’UTR isoforms. Furthermore, the same differences in distribution were observed under normoxia, indicating that the shorter 5’UTR isoforms are more efficiently translated than longer ones, regardless of oxygen tension (**Fig 7e).** Considering the limitation related to the primer design that did not allow full separation of the PDK1 5’UTR isoforms, we further confirmed these findings using dual luciferase reporter assays in HEK 293T cells (**Fig 7f**, raw values in **Fig S8f-g**). As expected^2,5,22^, hypoxia globally reduced translation driven by ACTB 5’UTR and all 5’UTR isoforms of PDK1, while translation of the ATF4 5’UTR reporter was sustained. However, under both hypoxia and normoxia, the reporters for the shorter (57 and 36 nt) isoforms were better translated than the longer (134 nt) isoform (**Fig 7f**). The same differences were observed under conditions of mTOR inhibition (INK128) and ISR activation (thapsigargin; TG) (**Fig 7g**, raw values in **Fig S8h-i**), further confirming that the hypoxia-enriched shorter 5’UTR isoforms of PDK1 mRNA are more efficiently translated both under normal and stress conditions. To further establish the impact of TSS switching on PDK1 mRNA translation, we expressed 134, 57, and 36 nt PDK1 5’UTR isoforms followed by CRISPR-mediated knockout of the endogenous PDK1 gene (**Fig S9a**). Cells expressing the 36 nt 5’UTR isoform produced more PDK1 protein relative to the amount of mRNA, both under hypoxia and normoxia (**Fig 7h-i, Fig S9b-c**), consistent with this isoform being more efficiently translated. To exclude the potential effects of hypoxia on the stability of PDK1 5’UTR mRNA isoforms, we performed actinomycin D chase, which did not reveal notable differences between 134 and 36 nt 5’UTR isoform stability under hypoxia (**Fig S9d-f**). This confirmed that induction of the shorter 5’UTR mRNA isoform of PDK1 under hypoxia was mediated by differential TSS usage and not by alterations in mRNA stability.

To examine the impact of TSS switching towards shorter 5’UTR isoforms of PDK1 under hypoxia, we subjected T47D cells expressing different PDK1 5’UTR isoforms to [^13^C] pyruvate labelling followed by stable isotope tracing analysis by GC-MS after 24 hrs in hypoxia (0.5% O_2_) or normoxia (20% O_2_). In cells expressing the short but not the long 5’UTR PDK1 isoforms, ^13^C-pyruvate tracing into lactate (lactate m+3) was increased upon exposure to hypoxia (**Fig 7j, Fig S9g-h**). As expected, ^13^C-pyruvate tracing into citrate (citrate m+2) was diminished across all lines grown under hypoxia relative to normoxia (**Fig S9i**, raw values in **Fig S9h,j**), with a modest reduction in conversion to alanine (alanine m+3) observed in hypoxia upon expression of the longer isoform (134 nt) (**Fig S9k,** raw values in **Fig S9h,l**). Taken together, these data suggest that cells expressing the shorter, more efficiently translated, 5’UTR isoform of PDK1 display enhanced lactate production under hypoxia.

Finally, we examined TSS switching for PDK1 after C48 treatment. Although TSS switching was not the same as under hypoxia, possibly due to the absence of strong transcriptional activation of PDK1 by HIF1, there was an enrichment of shorter, more efficiently translated 5’UTR isoforms (**Fig S9m-n**). Moreover, PDK1 protein levels were increased after C48 treatment although total mRNA levels were unchanged (**Fig S9o**), consistent with more efficient translation of the shorter 5’UTR isoforms. In addition, under hypoxia, expression of the shorter PDK1 5’UTR isoform was accompanied by extension of H3K4me3 around the TSS, a 25 base upstream shift in the dyad position of the +1 nucleosome, and an increase in occupancy at the upstream -2 nucleosome (**Fig 7k**). These changes were mirrored by C48 treatment, where H3K4me3 modestly extended around the TSS. NucleoATAC^52^ analysis of ATAC-seq (**Fig S9p**) on the same C48-treated cells revealed a similar upstream shift in the dyad of the +1 nucleosome by 36 bases, and an increase in occupancy at the upstream -2 nucleosome (**Fig 7k**) in absence of HIF1 induction (**Fig 4f**) or a change in PDK1 mRNA levels (**Fig S9o**). These findings suggest that modulation of H3K4me3 results in changes to nucleosome occupancy and positioning, together with altered selection of TSSs. Overall, this proposes a model where adaptive translational responses altering cellular phenotypes can be coordinated from the level of chromatin modifications altering TSS selection and remodelling 5’UTRome.

## DISCUSSION

In this study, we discovered that the microenvironmental stress hypoxia rewires gene expression via coordination of epigenetic, transcriptional, and translational programs. We uncovered widespread changes in TSS usage under hypoxia that were recapitulated by inhibition of the H3K4me3 eraser KDM5A and that were reversed by impeding MLL-associated COMPASS methyltransferase complexes. TSS switching modulated 5’UTRs of thousands of transcripts, altering potential for interactions with the translation initiation machinery. This hypoxia-associated 5’UTR pool occurs in parallel with reprogramming of the translational apparatus, and together contribute to a survival-enhancing adaptive translational program. Within this program, the switch towards glycolytic metabolism under hypoxia was facilitated by a shift in expression towards shorter, more efficiently translated 5’UTR isoforms of PDK1. Moreover, disruption of TSS switching led to a loss of cellular fitness under hypoxia, suggesting an important role during adaptation. Together, these findings support a mechanism of gene expression control whereby translational regulation is directed at the level of chromatin modifications, driving alternate TSS selection that gives rise to 5’UTR isoforms that exhibit different translation efficiencies (**Fig 8**). Over 50 years ago, it was proposed that mRNA translation may be selectively modulated by altering the availability of the components of the translation initiation machinery^57^. This can largely be explained by 5’UTR features that distinguish translationally “strong” mRNAs, that are efficiently recruited by the translation initiation apparatus and can thereby outcompete “weak” mRNAs. In this context, our results suggest that H3K4me3-dependent TSS selection under hypoxia may be adjusting the abundance of “strong” vs. “weak” 5’UTR isoforms to remodel the proteome in response to hypoxia.

**Figure 8:**
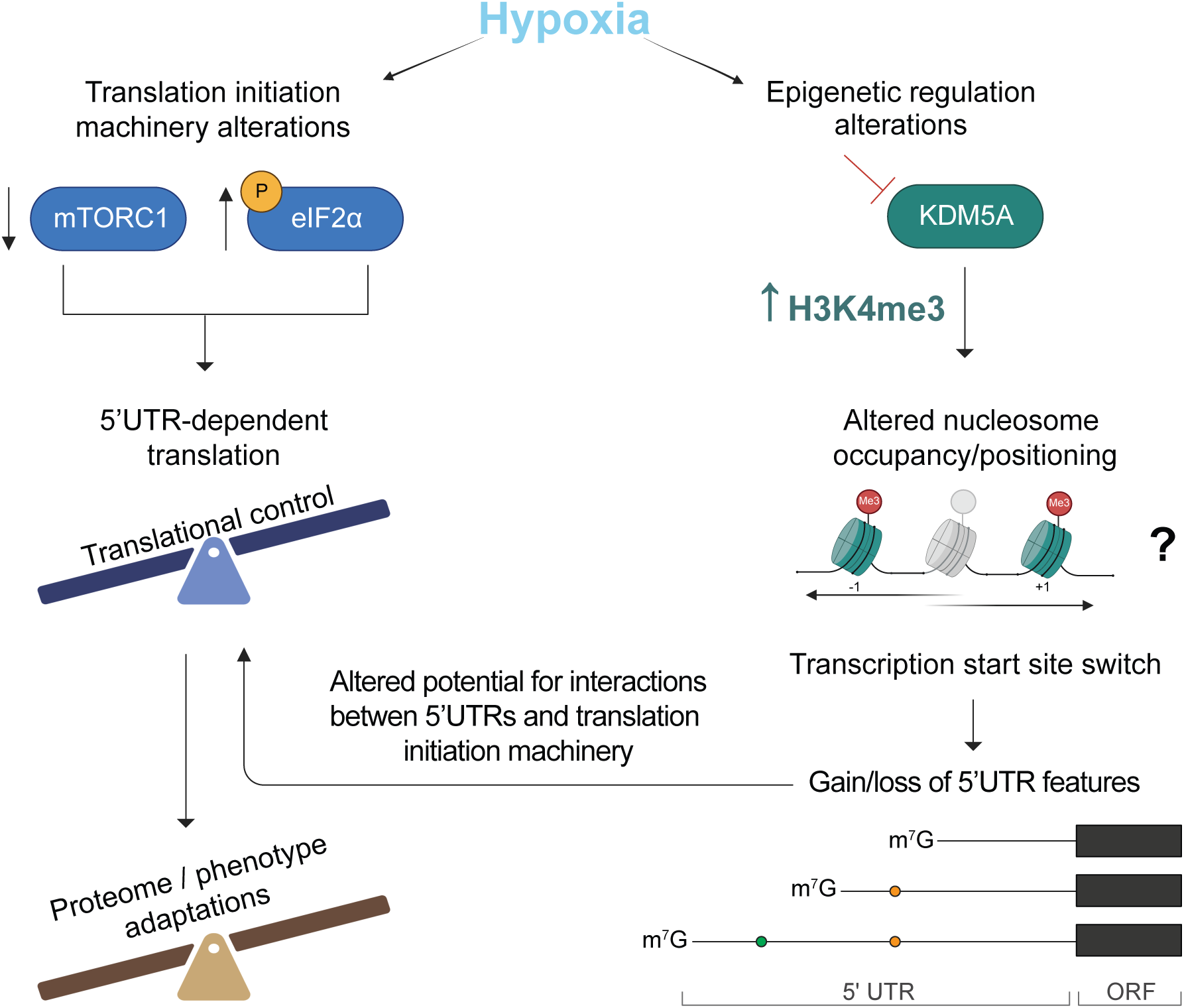
Epigenetic coordination of transcriptional and translational programs in hypoxia. Hypoxia induces reprogramming of the translational machinery largely through inhibition of mTORC1 and activation of the ISR via increased phosphorylation of eIF2 . These events together lead to global suppression of mRNA translation, and activation of transcript-selective translation that promotes proteome adaptations to hypoxia and associated phenotypes. Concurrently, hypoxia leads to remodelling of H3K4me3 due to loss of oxygen-dependent activity of KDM5A. This alteration of H3K4me3 around TSSs leads to changes in nucleosome occupancy and positioning and TSS switching for a subset of genes. This extensive TSS switching alters the composition of regulatory mRNA features in 5’UTRs, changing their potential for interactions with the translation initiation machinery. The epigenetically-mediated TSS switching produces a pool of 5’UTR isoforms, that in concert with the hypoxic translational machinery, help to drive changes in the proteome that are fundamental to cellular adaptations to hypoxia.

Most genes have multiple TSSs, with differential use linked to tissue-specificity, and disease states^58–60^. Our previous work identified several stem cell factors with 5’UTR isoforms with different translation efficiencies under hypoxia that arise from distinct TSSs^26^. In colon cancer cells, hypoxia was found to induce differential promoter usage for 191 genes^61^. Here, specifically focusing on TSS switching events that would not alter the resulting proteoform, and profiling changes within single promoters, we found that TSS switching events under hypoxia are dramatically more widespread than previously appreciated, affecting more than 20% of protein-coding transcripts across highly divergent cell types. The importance of this high-resolution approach is illustrated by 5’UTR regulatory elements that are highly positional, such as TOP motifs, where a difference of only a few nucleotides can drastically alter translation of corresponding mRNAs^25^. Indeed, we found that TSS switching leading to changes in TOP motifs plays a significant role in translational reprogramming of H9 cells in response to hypoxia. Furthermore, even modest changes in 5’UTR length were significantly associated with altered translation efficiency under hypoxia, demonstrating the surprising impact of this unappreciated regulatory mechanism.

Dynamic TSS switching resulting in differentially translated 5’UTR isoforms has also been reported to occur during meiosis and response to ER stress in yeast^39,40^. This mechanism has been shown to be driven by factors that initiate transcription from upstream TSS producing 5’UTRs containing inhibitory uORFs. Interestingly, our findings in human cells have also revealed extensive TSS switching leading to the gain or loss of uORFs and altered translation efficiency, suggesting that this mechanism may also play a role in mammalian cells responding to hypoxia. However, only 14% and 10% of the TSS switching in T47Ds and H9, respectively altered the presence of uORFs in the 5’UTR, indicating that the vast majority of the TSS switching in human cells under hypoxia impacts translation through distinct mechanisms.

While the role of mTOR inhibition and ISR activation in mediating global suppression of protein synthesis under hypoxia has been well established^5^, other mechanisms governing selective translation of survival-promoting transcripts remain less understood. Our study uncovers potential implications of other factors that may be orchestrated with epigenetic remodeling of the 5’UTRome, including U34 tRNA modifications^49^, the RNA helicase DHX9^41^, DAP5^42^, and alterations in eIF4E phosphorylation^45^. Interestingly, we also identified several previously unrecognized sequence motifs in 5’UTRs that are impacted by TSS switching and associated with both translational activation and suppression under hypoxia, warranting future studies.

A previous study in RCC4 renal cell carcinoma cells identified HIF-dependent changes in TSS usage, impacting translation^31^. We also report significant TSS switching at HIF1 targets, including PDK1. However, we found that the vast majority of TSS switching occurred for genes that are not known to be transcriptionally regulated by HIF1. Furthermore, some of these changes could be reproduced solely by modulating H3K4me3, in absence of HIF1 induction and independent of alterations in mTOR signaling or ISR. The precise role of H3K4me3 in gene expression has been debated^9,13^ but this mark has been shown to participate in anchoring transcriptional machinery to nucleosomes, suggesting a role in transcriptional activation^62^. Furthermore, local induction of H3K4me3 was sufficient to enable transcription at silenced loci in several cell models^63^. In contrast, loss of H3K4me3 has been found to have a limited effect on transcription, and it is not required for transcription to occur in most cases^9,13,64^. Here, we showed that inhibition of KDM5 leading to an increase and redistribution of H3K4me3 resulted in extensive TSS switching that partially reproduced changes observed under hypoxia, most often in the absence of changes in mRNA abundance. Conversely, inhibition of MLL containing COMPASS methyltransferases^65^ under hypoxia blocked a subset of TSS switching events from occurring. This suggests that H3K4me3 may play an important role in precise TSS selection under different cellular conditions, and modulation of H3K4me3 appears to be sufficient to alter TSS selection for a subset of genes. We also observed changes in nucleosome occupancy and positioning associated with TSS switching both under hypoxia and with KDM5A inhibition, suggesting a possible mechanism wherein modifications in H3K4me3 drive TSS selection concomitant with changes in nucleosome positioning and occupancy.

Although we focused on the orchestration of the epigenetic events with translation initiation under hypoxia, the elongation phase of translation may also be impacted via, e.g., modulation of eEF2K activity^66^ and/or changes in methylation of eEF1A^67^ thus warranting further studies. In addition, although we focused only on TSS switching that could be linked to gene-level changes in translation efficiency or alterations in the proteome, we also observed extensive TSS switching altering 5’UTR isoform levels for genes that appeared to be unregulated under hypoxia, suggesting a mechanism of isoform-level offsetting that may act to maintain constant protein levels. This observation raises interesting questions regarding the role of TSS switching in proteome homeostasis in the context of cellular stress, and warrants future studies.

In conclusion, we present a mechanism whereby translational adaptations under hypoxia are tuned by alterations in the availability of 5’UTR sequences that are coordinated by epigenetically-mediated changes at TSSs. These findings put forward a largely unappreciated paradigm that cellular stress can be sensed at the level of chromatin to direct adaptive remodeling of the translatome.

## METHODS

### Primer sequences

All primer sequences are provided in **Supplemental File S7.**

### Cell lines

T47D cells (ATCC) were grown in RPMI-1640 Medium (Gibco) with 10% fetal bovine serum (FBS) (Corning). 293T cells were grown in DMEM (Gibco) with 10% FBS. Cells were maintained using standard procedures. Cells were authenticated at the Sick Kids Research Institute and tested for mycoplasma (Mycoplasma detection kit, ATCC). H9 hESCs were obtained from WiCell, and the use of hESCs was approved by the Stem Cell Oversight Committee of Canada. hESCs were cultured on plates prepared with Corning® Matrigel® hESC-Qualified Matrix (Stemcell Technologies) and fed daily using pre-warmed mTeSR™1 feeder-free medium supplemented with mTeSR™1 5X Supplement (Stemcell Technologies). Colony quality, confluence, and spontaneous differentiation were visually monitored daily, and differentiated colonies were manually scraped off the plate. hESCs were passaged using StemPro™ EZPassage™ Disposable Stem Cell Passaging Tool (Thermo Fisher). All tissue cultures were maintained at 37 °C and 5% CO_2_ in a humidified environment.

### Plasmids

The 5’UTR sequences of beta-actin (NM_001101.5) and mouse ATF4 (NM_009716.3), and the sequences spanning 134 nt and 57 nt of the PDK1 5’UTR were synthesized as gBlocks (IDT) containing the T3 promoter and upstream *Nde*I and downstream *Mlu*I restriction sites. They were subcloned into the bicistronic luciferase reporter pKS-FF-HCV-Ren^68^ using a standard restriction digest/T4 DNA ligation protocol. The shorter 36 nt 5’UTR of PDK1 was synthesized as Ultramer DNA oligo containing the T3 promoter sequence and flanking sequences overlapping the digested vector. They were inserted into the digested pKS-FF-HCV-Ren plasmid using the NEBuilder HiFi DNA Assembly Cloning Kit (New England Biolabs).

The plasmid pGL4.13_134nt-PDK1 containing the 134 nt long 5’UTR and the complete ORF of PDK1 was constructed as follows. The 134 nt 5’UTR was amplified by PCR (primer pair PDK1-UTRstd_forw and PDK1_UTRstd_rev) from the pKS-134nt-PDK1-FF-HCV-Ren plasmid. The PDK1 open reading frame (ORF) was amplified from pDONR223-PDK1 (addgene #23804) (primer pair PDK1_ORFstd_forw and PDK1_ORFstd_rev). A pGL4.13 plasmid, modified by introduction of a *BsiW*I restriction site downstream of the SV40 promoter (pGL4.13_BsiWI-PacI), was digested with *BsiW*I (Pfl23II) and *Xba*I (Thermo Scientific) to excise the 5’UTR and luciferase ORF. The two PCR fragments containing the PDK1 5’UTR and the PDK1 ORF were inserted into the digested vector using the NEBuilder HiFi DNA Assembly Cloning Kit, creating the plasmid pGL4.13_134nt-PDK1.

The two PAM motifs within the PDK1 ORF to be targeted by sgRNAs/CRISPR were modified by site-directed mutagenesis using the Platinum SuperFi PCR mastermix and overlapping PCR primers (PDK1_PAMmut_forw and PDK1_PAMmut_rev) containing the mutation to create pGL4.13mod-134nt-PDK1-PAM. The pUltra plasmid (addgene #24129) was used as a backbone to create 5’UTR-isoform PDK1 rescue plasmids for use in lentiviral transduction. The pUltra plasmid was modified by cloning a second EGFP ORF in the position downstream of the P2A sequence to create pUltra_2xEGFP (primer pair EGFP_XbaI_forw and EGFP_XbaI_rev). The fragment containing the Ubc promoter and the first EGFP ORF was then excised by restriction digest with *Pac*I and *BsrG*I and replaced by a fragment amplified from pGL4.13mod-134nt-PDK1-PAM containing the SV40 promoter and PDK1 5’UTR and ORF and flanking *Pac*I and *BsrG*I restriction sites (primer pair SV40-PDK1-pUltra_F and SV40-PDK1-pUltra_R), creating the plasmid pUltra-SV40-134nt-PDK1-PAM. Plasmids containing the shorter 5’UTRs of 57 nt and 36 nt length were created by amplifying the fragment containing the respective 5’UTR, PDK1 ORF, P2A sequence and EGFP ORF flanked by upstream *BsiW*I and downstream *EcoR*I sites from pUltra-SV40-134nt-PDK1-PAM and subcloning it in the *BsiW*I and *EcoR*I digested plasmid pUltra-SV40-134nt-PDK1-PAM (primer pairs BsiWI_PDK1-57 or BsiWI_PDK1-36nt and pUltra_P2A-EcoRI_rev).

All cloning/propagation with the lentivirus pUltra backbone was performed in NEB® Stable Competent E. coli. One Shot™ MAX Efficiency™ DH5α-T1R Competent Cells were used for all other plasmids. All plasmids were confirmed by Sanger sequencing.

### PDK1 CRISPR knock-out and 5’UTR isoform rescue

The following procedure describes the generation of T47D-KO cells stably expressing PDK1 from transcripts containing 134 nt, 57 nt or 36 nt long PDK1 5’UTR followed by the complete PDK1 ORF and in frame P2A and eGFP. The derived cell lines are 134 nt (clones #4 and #9) 57 nt (clones #4 and #9) and 36 nt (clones #4 and #14). The cell line EV expresses PDK1 from the endogenous locus and eGFP from lentiviral overexpression.

Overexpression of 134 nt, 57 nt and 36 nt 5’ UTR isoforms of PDK1 was carried out using the plasmids pUltra-SV40-134nt-PDK1-PAM, pUltra-SV40-57nt-PDK1-PAM and pUltra-SV40-36nt-PDK1-PAM described above. These plasmids encode the various PDK1 5’UTRs, PDK1 ORF, P2A sequence and eGFP ORF. pUltra plasmid containing EGFP but no gene insert was utilized as a negative control to create T47D EV.

For lentiviral production, 2.5×10^5^ HEK293T cells were seeded in a 6 cm dish (Sarstedt) to achieve 40-50% confluency the following day. Cells were then co-transfected with 4 µg of respective PDK1 5’UTR isoform-containing or empty-pUltra plasmid, 2.66 µg of psPAX2 packaging plasmid (Addgene 35002), and 1.66 µg of pMD2.G plasmid (Addgene 12259) using the jetPRIME transfection reagent as described by the manufacturer’s protocol (Polyplus transfection). The growth media was changed 24 hrs later and collected 48 hrs post-transfection. Virus-containing media was filtered through a 0.45 µm filter (Frogga Bio) and was mixed with fresh media at a 1:1 ratio. Polybrene (Sigma-Aldrich) was added to obtain a final concentration of 8 µg/mL. The mixture was added to 1.0×10^4^ T47D cells that were seeded in a 6-well plate (BioBasic). Cells were re-transduced the following two days with a mixture containing virus media collected at 72 hrs and 96 hrs post-transfection, respectively. T47D cells were allowed to recover for 48 hrs after the last transduction. Enhanced green fluorescent protein (eGFP) was used as the selection marker. Fluorescence-activated cell sorting (FACS) was conducted using a BD FACSAria™ Fusion Flow Cytometer (BD Biosciences) and approximately 1.0×10^4^ T47D cells expressing respective 5’UTR PDK1 isoforms were sorted into separate 6-well dishes (BioBasic) as a population and later expanded.

In order to deplete endogenous PDK1 expression, two guide RNAs (gRNAs) targeting exon 2 of hPDK1 (against hPDK1 sequences GCAAGAGTTGCCTGTCAGACTGG and TTGCCGCAGAAACATAAATGAGG) were designed using the CHOPCHOP webtool^69^ and purchased with the appropriate overhangs in order to be cloned into the lentiCRISPRv2 plasmid conferring G418 resistance (Addgene 98292). Cloning was carried out as described in the Target Guide Sequence Cloning Protocol by Zhang *et al.*^70,71^. Modifications to the mentioned protocol included: utilizing BsmBI-v2 (New England Biolabs) to digest the lentiCRISPRv2 plasmid, using the Zymoclean Gel DNA Recovery Kit (Zymo Research), and carrying out the ligation reaction overnight at 4 °C. Transformation of the ligated plasmid was carried out via 42°C heat-shock into chemically competent Stbl3 cells (Fisher Scientific). Bacteria were plated on Lysogeny Broth (LB)-agar plates (Biobasic) containing ampicillin (1 µg/mL) (BioBasic) and grown bacterial colonies were picked and inoculated in Lysogeny Broth with ampicillin (100 μg/mL) overnight. Extraction of plasmid DNA was conducted using the QIAprep Spin Miniprep Kit (Qiagen) and samples were sent for Sanger sequencing using a primer for the human U6 promoter (GAGGGCCTATTTCCCATGATT). Correct insertion of the gRNA into the plasmid was verified before proceeding.

Lentivirus production and transduction were carried out as previously described, using the two lentiCRIPSRv2 plasmids with the inserted gRNAs targeting hPDK1, and a lentiCRISPRv2 plasmid with no sgRNA insert as a negative control. Previously sorted T47D cells overexpressing different 5’UTR PDK1 isoforms or empty pUltra plasmid were seeded into separate 6-well plates (1.0×10^4^/well). The cells were transduced on three consecutive days. The cells were allowed to recover for 48 hrs prior to being selected with G418 (500 μg/mL) (Bio Basic) for an additional four days. Afterwards, single-cell sorting was conducted using FACS into 96-well plates (Sarstedt) containing conditioned media (45% filtered cultured media, 45% fresh media, 10% FBS). Approximately two weeks later, the sorted clones were expanded into 24-well plates (BioBasic) and later into 6-well plates in order to generate clonal cell lines.

Depletion of endogenous PDK1 protein levels was initially confirmed by immunoblotting. Candidate clones with loss of endogenous PDK1 expression were kept for further validation. Moreover, the genomic DNA loci of candidate clones were sequenced and analyzed for insertion-deletion mutations (INDELs). Two sets of PCR primers (Set 1 Forward: AATGATAGCAGCAACGGGGT, Reverse: AGGGCACAAGCCCAAATACA, Set 2 Forward: GCTCAGACATGCTCAGGTTAC, Reverse: CCACAACTCCATCAAATTGCAAAC) were designed in the intronic regions flanking the CRISPR target sites. Extraction of genomic DNA for each clone was carried out using the PureLink Genomic DNA Mini Kit (Fisher Scientific). Two rounds of polymerase chain reaction (PCR) on the extracted genomic DNA (35 PCR cycles with Set 1 followed by 45 PCR cycles with Set 2) were carried out and gel electrophoresis was utilized to separate the PCR products. The smaller PCR product for each clone was excised and purified using the Zymoclean Gel DNA Recovery Kit (Zymo Research). The products were sent for Sanger sequencing along with the Set 2 forward and reverse primers. The chromatograms were analyzed for the presence of mutations causing a premature stop codon or a frameshift.

### PCR on genomic DNA

Genomic integration of the regions encoded by the lentiviral transfer plasmids pUltra-SV40-134nt-PDK1-PAM, pUltra-SV40-57nt-PDK1-PAM or pUltra-SV40-36nt-PDK1-PAM was confirmed by extraction of genomic DNA from the respective cell lines using the PureLink genomic DNA mini kit (Invitrogen) and PCR using the AmpliTaq 360 gold mastermix (Applied Biosciences #4398881) and primers spanning the region between the SV40 promoter and the beginning of the PDK1 ORF according to the manufacturer’s instructions. The size of the PCR product was determined by agarose gel electrophoresis.

### Cell culture experiments

For all experiments, T47D cells were plated 24 hrs prior to the treatment (1.2 × 10^6^ in 60 mm dishes or 5.8 ×10^5^ in 6-well plates). For hypoxia experiments, media was changed to fresh RPMI 10% FBS and cells were incubated in normoxia (regular incubator) or hypoxia (Biospherix OxyCycler model C42 or ProOx model C21 chambers housed in a regular incubator) at 0.5% O_2_ for the indicated time. For hypoxia time course experiments, all cells received fresh media at the start time. 48 hrs hypoxia samples were placed in the hypoxia chamber at the start of the time course. 0 hr hypoxia samples were incubated in normoxia for 48 hrs. All other time points were incubated in normoxia until shifting them to hypoxia at 6, 12 or 24 hrs before the 48 hr collection time point.

For compound 48 (C48) (Axon Medchem, #2809/batch1) treatments, media was replaced with fresh RPMI 10% FBS containing 10 µM C48 or the equivalent volume of DMSO as vehicle control. For the C48 dose-response experiments, media was replaced with a 1:10 dilution series starting at 10 µM C48 in RPMI 10% FBS. For C48 ChIP experiments cells were treated with DMSO for 48 hrs, or 24 hrs DMSO followed by 10 µM C48 for an additional 24 hrs.

For OICR-9429 (S7833, Selleckchem) and hypoxia co-treatments cells were treated with 25 µM OICR-9429 or an equivalent volume of DMSO as vehicle control. After 48 hrs media was replaced with fresh media containing 25 µM OICR-9429 or DMSO and cells were placed under hypoxia or normoxia for an additional 48 hrs before harvesting cells for RNA or protein lysates.

### Proliferation

T47D cells were plated 24 hrs prior to treatment. At 0 hrs media was changed to fresh RPMI 10% FBS with 25 µM OICR-9429 or equivalent volume of DMSO. At 48 hrs cells received fresh RPMI 10% FBS with 25 µM OICR-9429 or equivalent volume of DMSO and were placed in normoxia or hypoxia (0.5% O_2_) for an additional 48 hrs before cell counting.

### RNA extraction and quantitative RT-PCR

T47D cells were placed on ice and washed with ice-cold PBS. 1 mL of Trizol (Invitrogen) was added, and cells were incubated at room temperature for 5 mins. RNA was extracted as per the manufacturer’s instructions, and RNA pellets were resuspended in RNase and DNase-free H_2_O. RNA concentration was determined with a Nanodrop and samples were stored at -80 °C. Fractions from polysome gradients were extracted using 1 mL Trizol LS reagent (Invitrogen) according to the manufacturer’s instructions. 500 ng RNA was reverse transcribed using the SuperScript™ IV VILO™ Master Mix with ezDNase™ (Invitrogen) at 55°C extension temperature according to the manufacturer’s instructions. After the RT reaction, cDNA was diluted 5-fold with H_2_O and 1 µl per replicate was used in all following real-time quantitative PCR (qPCR) reactions. Primer/probes for LOX (Hs00942480_m1), PDK1 (Hs01561850_m1), eGFP (Mr04329676_mr) and RPLP0 were used for qPCR with TaqMan fast advanced mastermix (Thermo Scientific #4444963).

qPCRs specific for the PDK1 5’UTR isoforms were performed using the SYBR™ Green PCR Master Mix (Applied Biosystems). The primer pair PDK1_93/80nt amplifies all PDK1 transcripts with 5’UTRs longer than 78 nt, primer pair PDK1_57/51nt amplifies all transcripts with 5’UTRs longer than 45 nt, and primer pair PDK1_ex2 amplifies all PDK1 transcripts. Relative amounts were determined using the relative standard curve method with a 1:4 serial dilution using cDNA from a hypoxia-treated sample.

To determine the relative contribution of the three main clusters of PDK1 5’UTR isoforms (longer than 78nt, between 78 nt and 45 nt and shorter than 45 nt) to the total transcript level (**Fig 7d-e**), the qPCR setup was modified as follows. qPCRs were performed as described above, except that the relative standard curve method was performed on a 1:5 dilution series of a linearized plasmid containing the full 134 nt of the 5’UTR plus the open reading frame (pGL4.13_134nt-PDK1) starting with 1 pg/µl of plasmid. The result of the qPCR with primer pair PDK1_93/80nt represents the UTR isoforms longer than 78 nt. The result of the qPCR with primer pair PDK1_57/51nt represents the transcripts longer than 45 nt, i.e., also includes the transcripts longer than 78 nt. The result of the qPCR with primer pair PDK1_ex2 represents all transcripts. The relative amount of just the mid-length or the short cluster was calculated as follows: mid-length = qPCR PDK1_57/51nt – qPCR PDK1_93/81nt: short = qPCR PDK1_ex2 – qPCR PDK1_57/51nt. The relative contribution to the total transcript level was calculated as the ratio of the relative amount of each cluster versus the relative amount of total transcript.

All qPCR reactions were run in triplicates on a QuantStudio5 or Viia7 cycler and analyzed using the QuantStudioTM Design & Analysis software v1.5.2 or the QuantStudio Real-time PCR software v1.6.1.

### Stability of transcripts with different PDK1 5’UTRs

T47D-PDK1-KO_134nt-PDK1-OE, 57nt-PDK1-OE and 36nt-PDK1-OE, and T47D EV cells were plated at 8 × 10^5^ cells per dish. After 24 hrs media was changed to fresh RPMI 10% FBS and cells were incubated in normoxia or hypoxia (0.5% O_2_). Cells were either harvested for RNA extraction at 8 hrs or treated with 5 µg/ml actinomycin D (Sigma, A9415) at 8 hrs and further incubated in normoxia or hypoxia for an additional 16 hrs before harvesting for RNA extraction. RT-qPCR was performed with TaqMan probes for eGFP and RPLP0 as described above and quantified using a relative standard curve using a 1:5 dilution of cDNA from untreated normoxic T47D EV RNA. Stability (ratio of treated/untreated) of 5’UTR isoform transcripts was assessed relative to the stability of RPLP0 transcripts to control for clonal differences. Induction of mRNA decay by actinomycin D treatment was confirmed with qPCR for unstable transcripts RIPK2 (Hs01572684_m1) and SOX2 (Hs04234836_s1).

### RLM-RACE

RNA ligase-mediated rapid amplification of cDNA ends (RLM-RACE) was performed with the FirstChoice™ RLM-RACE Kit (Invitrogen) according to the manufacturer’s instructions. In short, 10 µg of RNA isolated from normoxia and hypoxia-treated T47D cells was treated with calf intestinal phosphatase (CIP) to remove the 5’-phosphate from all molecules that contain free 5’-phosphates. RNA was extracted with acidic phenol/chloroform to remove CIP. The RNA was then treated with tobacco acid pyrophosphatase (TAP) to remove the cap structure from the full-length mRNA, leaving a 5’-monophosphate. A synthetic 5’RACE RNA adapter was ligated to molecules containing a 5’-phosphate. 2 µl of the ligated RNA was reverse transcribed using the SuperScript™ IV VILO™ Master Mix (Invitrogen) at an extension temperature of 55 °C. 1 µl cDNA was used to perform nested PCR with forward primers specific for the 5’RACE adapter sequence (5’RACE_outer or 5’RACE_forw_inner) and reverse primers specific for PDK1 (PDK1_rev_outer, PDK1_rev_inner) and the AmpliTaq Gold™ 360 Master Mix (Applied Biosystems). The first round PCR cycling conditions were 95 °C for 10 min for polymerase activation, followed by 32 cycles of 95 °C for 30 sec denaturation, 59 °C for 30 sec annealing and 72 °C for 1 min extension, followed by 72°C for 7 min extension and 4 °C on hold. The second round PCR cycling conditions were 95 °C for 10 min for polymerase activation followed by 35 cycles of 95 °C for 30 sec denaturation, 58 °C for 30 sec annealing and 72 °C for 1 min extension, followed by 72 °C for 7 min extension and 4 °C on hold. PCR products were separated on a TAE/agarose gel, excised and extracted using a GE gel extraction kit. The purified PCR fragment was then inserted into the vector pCR4-TOPO using the TOPO™TA Cloning™ kit (Invitrogen) and One Shot™ MAX Efficiency™ DH5α-T1R Competent Cells (Invitrogen). Plasmids were extracted using the Qiagen Plasmid mini kit, and the sequence of the 5’ends of the PDK1 transcripts was determined by Sanger sequencing.

### Polysome profiles

T47D cells (8 × 10^6^ cells per condition) were seeded in 15 cm plates 24 hrs prior to the experiment. Media was changed to fresh RPMI 10% FBS before cells were exposed to normoxia or hypoxia (0.5% O_2_) for 24 hrs as described above. Polysome profiles were generated as previously described^72^. Cells were pre-treated with 100 μg/mL cycloheximide for 5 min and scraped in ice-cold, cycloheximide supplemented (100 μg/mL) PBS, using rubber scrapers. Cells were then pelleted in 15 mL conical tubes at 240 × g for 5 min at 4 °C, and lysed on ice in 500 μL of hypotonic lysis buffer (5 mM Tris–HCl, pH 7.5, 2.5 mM MgCl_2_, 1.5 mM KCl, 100 μg/mL cycloheximide, 1 mM DTT, 0.5% Triton, 0.5% sodium deoxycholate) for 15 min. Subsequently, lysates were cleared through a 15 min, 20 817 × g centrifugation at 4 °C. Cleared lysates were diluted with hypotonic lysis buffer to set their optical density (OD) at 260 nm to 10–20. Sucrose gradients (5–50%) were generated in polypropylene tubes (Beckman Coulter, 331372, 14 × 89 mm) using a gradient maker (Biocomp Gradient Master 108), according to the manufacturer’s instructions. A volume of 500 μL was removed from the top of the sucrose gradient to allow for sample loading. Lysates were then carefully layered over the top of linear sucrose gradients and centrifuged at 4 °C for 2 hrs at 222,228 × g (36,000 rpm) using an ultracentrifuge (Beckman Coulter, Optima XPN-80, rotor SW41Ti), while 10% of the sample was kept as input. Ultracentrifuged samples were fractionated into 2 mL tubes using a density gradient fractionation system (Brandel, BR-188-177), resulting in 12 fractions of 800 μL each. Input samples, as well as sucrose gradient fractions, were mixed with 1000 μL of TRIzol LS reagent (Invitrogen) and kept at −80 °C for subsequent RNA isolation.

### Immunoblot analysis

Cells were placed on ice, washed with ice-cold PBS, lysed in RIPA buffer containing 1x Halt™ Protease and Phosphatase Inhibitor (Thermo Scientific) and cleared by centrifugation at 17,000 x g. Lysates for histone immunoblots were also sonicated (one pulse, 50% amplitude) prior to centrifugation. Protein concentration was determined with a Micro BCA™ Protein Assay Kit (Thermo Scientific) as per the manufacturer’s instructions and read on a Spectramax ABS plus plate reader.

Equal amounts of protein were separated by polyacrylamide gel electrophoresis and transferred onto a 0.45 µm nitrocellulose membrane (BioRad). Membranes were blocked with Odyssey blocking buffer, incubated with primary antibody in Odyssey Antibody dilution buffer overnight at 4 °C, washed in TBST, incubated with secondary antibody (IRDye 800CW or IRDye 680RD) matching the primary antibody species for 1 hr at room temperature and analysed on a LI-COR Odyssey CLx imaging system. Intensity of the bands was quantified with ImageStudio software. All antibodies are listed in **Supplemental File S7.**

### In vitro transcription

pKS-FF-HCV-Ren plasmids containing the 5’UTR isoforms of PDK1 were linearized by *BamH*I (Thermo Scientific) digestion and purified by standard phenol/chloroform extraction. *In vitro* transcription and capping were carried out using a modified protocol from Steinberger *et al.*^68^. In short, 3 μg linearized plasmid was transcribed with 10 µl T3 RNA polymerase (New England Biolabs), 1× RNA polymerase buffer (New England Biolabs), 1 mM CTP, 1 mM ATP, 1 mM UTP, 0.2 mM GTP, 1 mM 3′-*O*-Me-^m7^GpppG (Anti-Reverse Cap Analog [ARCA]; NEB, S1411), 100 U RNase Inhibitor (New England Biolabs) in a volume of 100 μL for 3 hrs at 37 °C. 2.5 µL DNaseI (5U, New England Biolabs) was then added, and samples were incubated for an additional 30 min at 37 °C. Samples were purified with the Invitrogen MegaClear kit according to the manufacturer’s instructions. RNA was quantified using a Nanodrop, aliquoted and stored at −80 °C.

### In cell translation experiments

293T cells (1.6 × 10^5^ per well) were seeded in 24-well plates. After 24 hrs cells were washed with serum-free media and then a mix of 150 ng *in vitro* transcribed RNA and 1 µL DMRIE-C reagent (Invitrogen) in 200 µL Opti-MEM was added directly to the cells as per manufacturer’s instructions. After 1 hr incubation (37 °C, 5% CO_2_) 200 µL DMEM 10% FBS containing Thapsigargin or INK128 (Cell Signaling) was added to the cells to a final concentration of 400 nM or 50 nM, respectively. Media containing an equal volume of DMSO was used as vehicle control. After an additional 7 hr incubation time cells were washed with PBS and lysed using 100 μL Passive Lysis Buffer (Promega).

For translation experiments under hypoxia, 293T cells (2×10^4^) were seeded in 24-well plates. After 24 hrs media was changed to fresh DMEM 10% FBS and cells were exposed to normoxia or hypoxia (0.5% O_2_) for 24 hrs as described above. Cells were then washed with serum-free media and a mix of 150 ng *in vitro* transcribed RNA and 1 µL DMRIE-C reagent (Invitrogen) in 200 µl Opti-MEM was added directly to the cells. After 1 hr incubation in normoxia or hypoxia 200 µL DMEM 10% FBS was added to the cells. After additional 7 hr incubation time in normoxia or hypoxia cells were washed with PBS and lysed using 100 μL Passive Lysis Buffer.

20 µL lysate was analyzed with the Dual Luciferase kit (Promega) as per the manufacturer’s instructions on a GloMaxO Navigator Luminometer. Transfections were performed in duplicates and each sample was then analysed in duplicate.

### Native ChIP-seq

Chromatin immunoprecipitation (ChIP) was used to analyze histone modifications. This protocol is based on previously published protocols from Shah *et al.*^73^. Three biological replicates were obtained for hypoxia and normoxia-treated T47D and H9 cells, and four biological replicates for C48 and DMSO-treated T47D cells. Post-treatment, control and treated cells were washed twice in PBS and collected (500 rcf for 5 min at 4 °C). Cell pellet volume was assessed before being flash frozen. Prior to each ChIP, antibodies were conjugated overnight at 4 °C in ChIP Buffer 1 (25 mM Tris pH 7.5, 5 mM MgCl_2_, 100 mM KCl, 10% glycerol, 0.1% NP-40, 200 μM PMSF, 50 μg/mL BSA) with 50/50 mix of Dynabeads™ Protein A (Thermo Fisher) and Dynabeads™ Protein G (Thermo Fisher) at a ratio of 6 μg of antibody to 25 μL of beads per sample. Prior to IP, beads were washed twice with ChIP Buffer 1 and resuspended in 25 μL of ChIP Buffer 1. Frozen cells were washed twice in Buffer N (15 mM Tris Base, 15 mM NaCl, 60 mM KCl, 8.5% sucrose, 5 mM MgCl_2_, 1 mM CaCl_2_, 1 mM DTT, 200 μM PMSF, 1x protease inhibitor, 50 μg/mL BSA), then resuspended in 2X cell pellet volume of Buffer N and lysed for 10 min on ice in an equivalent volume of 2X Lysis Buffer (1X Buffer N, 0.6% NP-40). Nuclei were pelleted (500 rcf for 5 min at 4 °C) and resuspended in 6X cell pellet volume of Buffer N, then spun through a Sucrose Cushion (15 mM Tris Base, 15 mM NaCl, 60 mM KCl, 30% sucrose, 5 mM MgCl_2_, 1 mM CaCl_2_, 1 mM DTT, 200 μM PMSF, 1x protease inhibitor, 50 μg/mL BSA) at 500 rcf for 12 min at 4 °C. The nuclei pellet was resuspended in 2X cell pellet volume of Buffer N. 2 μL of nuclei suspension was diluted with 18 μL of 2 M NaCl in triplicate, vortexed, and sonicated with a water bath sonicator (30X cycle of 30 sec on, 30 sec off) prior to measuring the chromatin concentration with a NanoDrop Spectrophotometer (Thermo Fisher). The concentration of nuclei suspension was then adjusted to 1 μg/μL and separated into 100 μg chromatin aliquots. In the case of excess nuclei, unused aliquots were flash-frozen.

Nuclei suspensions were prewarmed for 2 min at 37 °C, then 46 U (2 µl) of MNase (Worthington Biochemical Corporation) was added for 12 min at 37 °C on a ThermoMixer (Eppendorf) set to 900 rpm. Digestion was stopped by adding 1/10 volume of MNase Stop Buffer (100 mM EDTA and 10 mM EGTA) and placing nuclei on ice. 5 M NaCl was slowly added to the suspension to a concentration of 0.6 M on the slowest vortex setting to release the chromatin, insoluble nuclei were then pelleted (18,000 rcf, 1 min, 4 °C). 15 μL of the soluble chromatin was saved for visualization and the rest of the supernatant was added to 0.07 g of hydroxyapatite (HAP) resin (Bio-Rad, Cat#157-0021) rehydrated in HAP Buffer 1 (5 mM NaPO4 pH 7.2, 600 mM NaCl, 1 mM EDTA, 200 μM PMSF) and rotated for 10 min at 4 °C. HAP mixture was run through Ultrafree-MC Centrifugal Filter (Millipore) at 600 rcf for 30 sec at 4 °C. Then the column was washed 4X with 200 μL HAP Buffer 1, 4X with 200 μL HAP Buffer 2 (5 mM NaPO_4_ pH 7.2, 100 mM NaCl, 1 mM EDTA, 200 μM PMSF), and eluted 3X with 100 μL of HAP Elution Buffer (500 mM NaPO_4_ pH 7.2, 100 mM NaCl, 1 mM EDTA, 200 μM PMSF). HAP input and eluates were visualized using 2% agarose gel in 1X TBE to confirm fragment sizes. Chromatin concentration of the combined HAP eluate was measured in triplicate using NanoDrop. Final chromatin concentration was adjusted to 20 ng/μL using ChIP Buffer 1 prior to IP.

15 μL of chromatin was set aside for input control and chromatin was added to beads conjugated with H3K4me3 antibody (Epigentek, A4033). Chromatin and bead mixture was rotated for 10 min at 4 °C, washed twice with ChIP Buffer 2 (25 mM Tris pH 7.5, 5 mM MgCl_2_, 300 mM KCl, 10% glycerol, 0.1% NP-40, 200 μM PMSF, 50 μg/mL BSA), washed once with ChIP Buffer 3 (10 mM Tris pH 7.5, 250 mM LiCl, 1 mM EDTA, 0.5% Sodium Deoxycholate, 0.5% NP-40, 200 μM PMSF, 50 μg/mL BSA), and washed once with TE buffer, transferring beads to a new tube after each wash step. DNA was finally eluted in 50 μL of ChIP Elution Buffer (50 mM Tris pH 7.5, 1 mM EDTA, 1% SDS) at 600 rpm for 10 min at 55 °C on a ThermoMixer. 35 μL of ChIP Elution Buffer was also added to the input control to bring the final volume to 50 μL. To both input and eluted ChIP DNA, 2 μL of 5 M NaCl, 1 μL of 500 mM EDTA, and 1 μL of 20 mg/mL Proteinase K was added and incubated for 2 hrs at 55 °C and DNA was purified using MinElute PCR Purification Kit (Qiagen). DNA was eluted in ultrapure H_2_O and concentration was measured before and after purification using Qubit dsDNA Quantitation, High Sensitivity (Thermo Fisher).

ChIP DNA was sent to the McGill University and Genome Quebec Innovation Center or the Institute for Immunology and Cancer (IRIC, Montreal), where it was further validated via Bioanalyzer, followed by shotgun library preparation, and library sequencing via Illumina NovaSeq6000. Samples were sequenced to a depth between ∼50-80 and ∼75-250 million reads per sample for H3K4me3 and inputs, respectively.

### Analysis of H3K4me3 ChIP-seq data

Raw H3K4me3 fastq files were processed using the nf-core/chipseq pipeline v.1.2.0 (available at https://github.com/nf-core/chipseq)^74^. Reads were aligned to GRCh38/hg38 in paired-end mode and all other parameters were kept as default. The output of the pipeline includes BAM files with aligned reads with duplicates, multi-mapping, unpaired, and ENCODE blacklist^75^ reads filtered out. In order to assess shifts in the position of H3K4me3 marks relative to TSSs of protein-coding genes, the data was converted to a transcript-rather than genome-centric format to facilitate comparisons to TSSs mapped using nanoCAGE sequencing. To do this, the filtered aligned reads were extracted from the BAM files and re-aligned to a custom index of 5’UTR genomic sequences using Bowtie (version 1.2.2)^76^ (settings: -v 2 -X 1000 -a). This custom index was created from genomic sequence regions corresponding to RefSeq transcripts (release 109, 2020-11-20)^77^, with 2,000 bp of sequence added upstream of the annotated TSS, and 10,000 bp added downstream of the TSS. BEDtools^78^ genomeCoverageBed (v2.29.1) was then used to compute the genome coverage of H3K4me3 with -dz and -pc options, and - scale to normalize for library size. The cumulative sum of H3K4me3 coverage was calculated for each genomic region corresponding to RefSeq transcripts, and genomic regions were trimmed at the nucleotide position where 95% of the cumulative H3K4me3 signal had already occurred. This was found to be necessary as H3K4me3 signals had variable degrees of extension around TSSs, such that the selected 12,000 bp window was not appropriate for all genes. In order to reduce file sizes and computation time, the position (relative to the TSS), and the H3K4me3 coverage were collected for every 0.1% increase in the cumulative sum of H3K4me3 signal, such that each genomic region corresponding to RefSeq transcripts would have 1,000 data points summarizing the distribution of the histone mark. Genomic regions corresponding to RefSeq transcripts were filtered out if they had a maximum scaled read coverage below 0.8, or if the background signal (from input control samples) was above 0.6.

To assess whether there were significant changes in the distribution of H3K4me3 around the TSSs for protein-coding genes, the data were filtered to only include transcript IDs that were detected in the nanoCAGE datasets for T47D and H9 cells. To detect directional shifts in the position of H3K4me3 signal around TSSs between hypoxia and normoxia, a two-sided Wilcoxon Mann-Whitney test was applied to the distributions of positions where 0.1% increases in the cumulative H3K4me3 signal had occurred for each condition. The directionality of the H3K4me3 shifts (upstream or downstream), was determined by the difference between the positions of the distributions at the 25^th^, 50^th^, and 75^th^ quantiles. To identify cases where a change in the distribution of the H3K4me3 mark had occurred, but without a defined directionality (e.g., a broadening of the peak, that extends both upstream and downstream), a two-sided Kolmogorov-Smirnov test was applied. *P*-values were adjusted for multiple testing using the Benjamini-Hochberg method, with a significance threshold of FDR < 0.01. As the biological replicates in both cell lines were reproducible (**Fig S6a-b**), the mean of the position distributions across the 3 replicates was used in statistical tests. Empirical cumulative distribution function, and scaled coverage plots show the mean of the replicates, with the standard deviation from the three biological replicates shaded.

### Omni-ATAC-seq

ATAC-seq libraries were generated following a modified Omni-ATAC-seq protocol^79^. Following treatment, both control and treated samples were washed, trypsinized and counted in a hemocytometer. To check viability, cells were counted with 0.4% trypan blue. Cell viability > 85% is required. After counting, 100,000 cells were pelleted at 500 rcf for 5 min at 4 °C and lysed on ice for 3 min with ATAC-Resuspension Buffer (10 mM Tris-HCl pH 7.4, 10 mM NaCl and 3 mM MgCl2) containing 0.1% Tween-20 (Fisher cat no. BP337), 0.1% IGEPAL® CA-630 (Sigma cat no. I3021), 0.01% Digitonin (ThermoFisher Scientific cat no. BN2006). After incubation with lysis buffer, cell lysates were washed with ATAC-Resuspension Buffer containing 0.1% Tween-20. After washing, nuclei were pelleted at 500 rcf for 10 min at 4 °C. Pelleted nuclei were resuspended in transposition mix containing the Illumina Tagment DNA TDE1 Enzyme (Tn5) and Buffer (Illumina cat no. 20034197), 0.1% Tween-20 and 0.01% Digitonin, and incubated in a thermomixer at 1000 RPM for 30 min at 37 °C. After tagmentation, fragments were cleaned using the MinElute PCR Purification Kit (Qiagen cat no. 28004). Cleaned-up fragments were resuspended in ultrapure water. The transposed fragments were then amplified with NEBNext High-Fidelity 2X PCR Master Mix (New England BioLabs cat no. M0541S) and indexed with Illumina Nextera XT Index Kit (Illumina cat no. FC-131-1001). Final libraries were cleaned-up and size-selected with AMPure XP SPRI Reagent (Beckman Coulter cat no. A63880). To assess library quality, 1 μL of each sample was submitted to Agilent Bioanalyzer at Queen’s University Department of Pathology and Molecular Medicine. Finally, samples were sent to Genome Quebec Innovation Center and sequenced on an Illumina NovaSeq6000 platform using a 100 base-pair paired-end read set-up.

### Analysis of ATAC-seq data

Raw ATAC-seq fastq files were processed using the nf-core/atacseq pipeline 2.1.2 (available at https://github.com/nf-core/atacseq)^74^. Reads were aligned to GRCh38/hg38 in paired-end mode and all other parameters were kept as default. As for ChIP-seq data, duplicates, multi-mapping, unpaired, and ENCODE blacklist^75^ reads were filtered out of final BAM files. BAM files from biological replicates were then merged, and consensus peak regions from MAC2 were extended by 1000 bp on each side using the BEDtools^78^ slop function (v2.29.1). Merged BAM files and extended consensus peaks were then processed using the NucleoATAC^52^ run function (v0.3.4) with default settings. Smoothed NucleoATAC signal outputs, and nucleosome dyad position BED files were visualized using Integrative Genomics Viewer (IGV)^80^. Changes in nucleosome position and occupancy for genes undergoing TSS switching were assessed using DeepTools (v3.4.3) computeMatrix with the –referencePoint “TSS” option, anchoring around the most abundance TSS peak detected by nanoCAGE sequencing under normoxia. Nucleosome occupancy and position around TSSs were visualized and significant differences between signal curves were detected using deepStats (v0.3.1) dsCompareCurves (available at https://github.com/gtrichard/deepStats)^81^.

### NanoCAGE library preparation, sequencing, and data preprocessing

The quantity and quality of total RNA from hypoxia-treated T47D and H9 cells, hypoxia and OICR-9429 or DMSO-treated T47D cells, and C48 or DMSO-treated T47D cells were assessed by Qubit (Life Technologies) and Bioanalyzer (Agilent Technologies), and 75 ng of RNA was used as input material to prepare nanoCAGE libraries as described previously^23,49^ (hypoxia and normoxia: T47D n = 2/3 [one normoxia sample failed in sequencing], H9 n = 4; C48 and DMSO n = 4; OICR-9429 and DMSO in hypoxia n = 3). Note that the Template Switching Oligonucleotides (TSO) include 8 nt random unique molecular identifiers (UMIs) that allow for identification of duplicate transcripts arising from PCR amplification, and one of six barcodes for increased multiplexing (ACAGAT, GTATGA, ATCGTG, GAGTGA, TATAGC, GCTGCA). Sequencing was performed at the SciLifeLab NGI (Sweden) facility. Hypoxia and normoxia libraries were sequenced using the HiSeq2500 (HiSeq Control Software 2.2.58/RTA 1.18.64) with a 100-base pair single-end setup using HiSeq Rapid SBS Kit vs chemistry. Bcl to fastQ conversion was carried out using bcl2fastq_v2.19 from the CASAVA software suite. The quality scale was Sanger/phred33/Illumina 1.8+. Libraries from C48 and OICR-9429 experiments were sequenced using the NovaSeq6000 using the same read set-up, and Bcl to fastQ version was carried out using bcl2fastq v2.20.0.422.

NanoCAGE sequencing reads with each of the six barcodes were extracted using TagDust (version 2.33)^82^, and Nextera XT N-Series 3’ adapters were trimmed using Cutadapt (v1.18)^83^ (settings: -e 0.15 -01 -n 4 -m 25). PCR duplicates were defined as reads with identical UMI and sequence in the first 25 bp, and were filtered to keep only one unique read. We have found this step to be essential for the quality of the downstream analysis. Ribosomal RNA was removed using bbduk.sh from the BBTools software suite, with default settings. A summary of the nanoCAGE preprocessing can be seen in **Fig S1a**. Next, reads were aligned to the human genome hg38 using Bowtie (v1.2.2)^76^ (settings: -a -m 1 --best --strata -n 2 -l 28), and reads that uniquely aligned or failed to align were collected for a second alignment step to a custom index of 5’UTR sequences. This custom index was created by extending RefSeq 5’UTR sequences (release 109, 2020-11-20)^77^ by 78 bases upstream using genomic sequence, and 78 bases downstream using mRNA sequences. These extensions were selected based on the criteria that the nanoCAGE read architecture allows for a maximal length of 78 after adapter trimming. Alignment to the custom index was also performed using Bowtie (settings: -a --best –strata -n 3 -l 25). Reads were trimmed by 1-2 based at the 5’ end in the event of alignment mismatches at the first or second positions, and strand invasion artifacts were removed using the Perl script provided by Tang *et al.*^84^ (settings: -e 2).

### Analysis of differential TSS usage

Transcription start site peaks were defined as the 5’ end mapping positions of reads for each sample. For hypoxia and normoxia conditions, peaks that were observed in less than n - 1 samples per condition were excluded to select only those that were reproducible. As the number of peaks detected was substantially higher for C48 and OICR-9429 experiments, only peaks observed in all samples (per condition) were retained. RefSeq transcripts with fewer than 10 reads were excluded before assessing library complexity by sampling increasing numbers of reads (in increments of 100,000) and counting the number of unique peaks (> 1 read) and RefSeq transcripts (> 10 reads) detected (**Fig S1b**). Reads corresponding to TSS peaks were then normalized to correct for library size (TPM, tags per million), and peaks with low expression (< 25% of the mean TPM expression per transcript) were filtered out. After normalization and filtering of non-reproducible and low-expression peaks, TSS clusters were defined using a dynamic sliding window approach with a window size of 5 nt. The beginning of a TSS cluster was defined as the position where the first TSS peak was encountered, and the cluster was extended until there were no longer TSS peaks within the span of the window. As differences in 5’UTR length may be more impactful for short 5’UTR isoforms than for long ones (e.g. a 30 nt vs. a 45 nt 5’UTR may have quite different translation efficiencies due to the difference in length, but the difference between a 215 nt vs. a 200 nt 5’UTR may not have similar implications), the size of the sliding window increased as the length of the 5’UTR increased, being scaled to be 2.5% of the length of the longest isoform for 5’UTRs longer than 200 nt. After TSS clusters were defined in each condition, overlapping clusters were unified between conditions to allow differential analysis of 5’UTR isoform expression. Once clusters were defined, reproducible and expression-filtered TSS peaks within cluster regions were quantified for each sample to generate a counts table for further analysis.

To assess differential expression of 5’UTR isoforms detected in both conditions (“quantitative” differences) and all replicates, TSS cluster counts were normalized using the “voom” function from the limma R package^85^. Significant changes in the relative expression of 5’UTRs isoforms between conditions were then identified using per-gene linear models with the design:

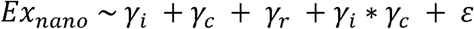

where ***Ex_nano_*** is the expression for each 5’UTR isoform across all conditions and replicates, *γ_i_* denotes the relationship with 5’UTR isoform, *γ_C_* is the relationship to the experimental condition, *γ_r_* is the relationship to replicate, *γ_i_* * *γ_C_* is the interaction between *γ_i_* and *γ*, and **ε** is the residual error. Similar to anota2seq^36^, a random variance model was then applied^86^. A low p-value for the *γ* * *γ_C_* interaction term indicates a significant difference in 5’UTR isoform expression depending on condition, that is independent of changes in overall transcript expression or replicate effects. When 5’UTR isoforms were not detected in all samples, the regression-based approach used for detecting the quantitative differences was not appropriate. Instead, a “qualitative” approach was used. For 5’UTR isoforms that were not detected in all samples, but were detected in at least two replicates in both conditions, a Fisher’s exact test was used to determine if there was a significant difference in expression of the 5’UTR isoform relative to all other isoforms between conditions (using the sums of nanocage sequencing reads per condition across all replicates). Finally, for a 5’UTR isoform that was only detected in one condition, Fisher’s exact test is not applicable, and significance was assessed using probabilities. In the condition expressing the unique 5’UTR isoform the probability of expressing the shared isoforms relative to all isoforms was first calculated. In the condition not expressing the unique isoform, the probability of expressing only the shared isoform was then calculated:

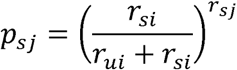

where **r** is the number of nanoCAGE sequencing reads summed across all replicates, **s** denotes shared isoforms, **u** denotes the uniquely expressed isoform, **i** the condition expressing the unique isoform and **j** the condition only expressing shared isoforms (i.e., *r_uj_* = 0).

All *p*-values from the different arms of the statistical analysis were collected and adjusted for multiple testing using the Benjamini-Hochberg method. Transcripts that underwent significant TSS switching under hypoxia treatment were considered to be those with an FDR < 0.15.

### Change-point analysis for Identification of enriched or depleted 5’UTR sequences resulting from TSS switching

After identifying transcripts that underwent significant TSS switching using the approach described above, we next determined more precisely which portions of the 5’UTR sequences were specifically enriched or depleted as a result of the TSS switching. For each transcript identified as undergoing significant TSS switching under the various treatment conditions, data from all biological replicates in each condition were pooled and the cumulative percentage of TPM-normalized reads was calculated at each TSS position. The cumulative percentages per TSS position under control (normoxic or DMSO) conditions were then subtracted from those under treatment (hypoxic or C48) conditions to provide the difference in cumulative percentage of reads at each TSS position. Next, a change-point algorithm was applied to identify the positions in the difference distributions where a change had occurred. This was implemented using the “cpt.mean” function from the changepoint R package^29^ with the arguments penalty = “AIC”, method = “SegNeigh”, and Q = 3 to allow up to two change-points to be identified. The longest 5’UTR isoform sequence for each transcript was then divided at the nucleotide positions defined by the change-points to generate three sequence segments based on their enrichment or depletion as a result of TSS switching. The enrichment or depletion of a 5’UTR sequence segment was quantified as the difference in the proportion of total TSS peak expression between conditions. Transcripts were also assigned a metric we termed the “TSS switch score”, defined as the maximum difference in expression across 5’UTR sequence segments. Higher TSS switch scores are indicative of TSS switching resulting in a greater change in the availability of particular 5’UTR sequences under a given treatment condition.

For transcripts with significant TSS switching, the percentage of GC content of 5’UTR sequence segments identified as enriched or depleted under hypoxia by change-point analysis was calculated and visualized using empirical cumulative distribution functions. Differences in GC content between hypoxia-enriched or depleted sequences were assessed using a Wilcoxon-Mann-Whitney test.

### Smart-seq2 library preparation, sequencing, and data preprocessing

RNAseq libraries for total and polysome-associated mRNA fractions from T47D and H9 cells cultured under hypoxia (1% O_2_) and normoxia (20% O_2_) (n = 4) were prepared using the Smart-seq2 protocol as previously described^87^. RNA quantity and quality were assessed by Qubit (Life Technologies) and Bioanalyzer (Agilent Technologies), and 10 ng of RNA was used as input material for each sample. After pre-amplification PCR 70 pg of cDNA underwent tagmentation in a total volume of 20 μL using the Nextera XT Kit (Illumina). Sequencing was then performed at the SciLifeLab National Genomics Infrastructure (NGI, Sweden) facility using the NovaSeq6000 (NovaSeq Control Software 1.6.0/RTA v3.4.4) with a 50-base paired-end setup using the ‘NovaSeqStandard’ workflow and S1 flow cell. Bcl to fastQ conversion was carried out using bcl2fastq_v2.20.0.422 from the CASAVA software suite. The quality scale was Sanger/phred33/Illumina 1.8+.

Adapters were removed from RNAseq reads using bbduk.sh from the BBTools software suite (sourceforge.net/projects/bbmap/) (settings k = 13 ktrim = n useshortkmers = t mink = 5 qtrim = t trimq = 10 minlength = 25). Ribosomal RNA was also removed using bbduk.sh, with default settings. Reads were aligned to hg38 using HISAT2^88^ (settings -- no-mixed --no-discordant), and uniquely mapping reads were quantified using the featureCounts function from the RSubreads package^89^ with RefSeq gene definitions and default settings.

### Standard RNA-seq library preparation, sequencing, and data preprocessing

RNA-seq libraries for C48 (10 μM, 24 hrs) and DMSO-treated T47D cells (n = 4) were prepared from total RNA by the Institute for Research in Immunology and Cancer (IRIC) Genomics Platform at the University of Montreal (Canada). Sequencing was performed on the Illumina Nextseq500 with a 75-base single-end setup. Data preprocessing was performed as described above for Smart-seq2 libraries, accounting for the single-end read setup.

### Analysis of differential translation

Raw counts from protein-coding genes with at least one read across all samples were TMM normalized^90^ and log_2_ transformed using the “voom” function^85^ as implemented in the anota2seq R package^36^. The reproducibility of data from hypoxia and normoxia-treated, and polysome-associated and cytosolic mRNA samples was assessed using Principal Component Analysis after filtering genes by variance to select those within the highest quartile of standard deviation across samples.

Changes in polysome-associated and cytosolic mRNA under hypoxia were analyzed using the anota2seq algorithm^36^ with the normalized count data. Batch effects were accounted for in the models by including sample replicates using the “batchVec” argument in Anota2seq. Default threshold settings were used [minSlopeTranslation = - 1, maxSlopeTranslation = 2, minSlopeBuffering = -2, maxSlopeBuffering = 1, deltaPT = deltaP = deltaTP = deltaT = log2(1.2)] with a significance threshold maxRvmPAdj = 0.15 (corresponding to an FDR < 0.15). Transcripts were classified into three modes of regulation (changes in mRNA abundance, changes in translation efficiency, and translational offsetting) using the “anota2seqRegModes’ function.

### Translatome modeling with Post-Transcriptional Network Modeling (postNet)

The translatome modeling performed required two inputs; the per-gene fold changes in translation efficiency or offsetting between hypoxia and normoxia that were found to be significant using anota2seq (described above), and a list of signatures describing regulatory variables for each gene (**Supplemental File S4**). The postNet ‘featureIntegration’ function was used with arguments: regOnly = TRUE, regulationGen = ‘translation’ or ‘buffering’, allFeat = TRUE, useCorel = TRUE, analysis_type = ‘lm’, covarFilt = 8, NetModelSel = ‘Omnibus’. Briefly, the software performs 3 steps. First, univariate linear models are used to identify significant associations between translational efficiency and individual variables. In the second phase, a stepwise regression is performed beginning with the variable that best explained changes in translational efficiency from step 1. Variables are added to the model iteratively, discarding those that do not improve the ability of the model to explain additional variance in translational efficiencies. This step both ranks the variables by importance and reveals covariance between them. In the final step, the covariance is removed to assess the adjusted contribution of each variable to the translational changes. This also allows calculation of the percent of variance in translation or offsetting explained by a given variable, adjusted for all other variables.

For the analysis presented in **Figs 3b-c and S3a-b**, and **Fig S4a-b** and **S5a-b,** for each RefSeq transcript the total number of nanoCAGE counts was summarized and the 5’UTR sequence corresponding to the mostly highly expressed position detected in the hypoxia condition was extracted. When multiple transcript isoforms were present for the same gene, the most highly expressed isoform was selected (i.e. based on the total number of nanoCAGE counts). If the expression of two or more isoforms was equal, the longest 5’UTR sequence was used. These sequences were then used to define sets of mRNA feature variables that may be able to explain the translational regulation observed. These included characteristics such as the GC content, length, and the presence of uORFs. Folding energy was also calculated using the Mfold algorithm^91^, and *de novo* 5’UTR motifs over-represented in regulated transcripts under hypoxia were identified using STREME^92^. G-quadruplexes were predicted using pqsfinder^93^ (v1.10.1) with a min_score = 47, and TOP motifs were quantified using the TOPscore method described by Philippe *et al.*^25^ using all reproducibly detected 5’UTR isoforms for a given transcript in the normoxia condition. The presence of an additional set of evidence-supported RNA binding protein (RBP) motifs obtained from the ATtRACT database^94^ was also quantified in 5’UTRs of transcripts that were regulated by translation or offsetting. Motifs were selected for inclusion in the final modeling using the ‘featureIntegration’ function of postNet (contrastSel = 1, regOnly = T, regulationGen = ‘translation’ or ‘buffering’, allFeat = TRUE, analysis_type = ‘lm’) to perform the same stepwise regression modeling described above, but only using the RBP motifs as variables to explain changes in translation or offsetting. Motifs that were significant in independently explaining proportions of the variance in translational changes were included in the final variable list (**Supplemental File S4**).

For the analysis presented in **Fig 3d-i and S3c-d,** and **Figs S4c-f** and **S5c-d** additional variables describing how 5’UTR features change as a result of TSS switching were also added to the models. These included the change in weighted mean 5’UTR length (as both the actual difference in log_2_ length, and as categorical values i.e., no change, shorter, longer), the difference in weighted mean GC content and folding energy, and the change in TOPscore between transcripts in hypoxia and normoxia. Finally, the TSS switch scores for each transcript from the change-point analysis were also added to the list of variables.

All of the translatome models also included known signatures of translational regulation by various genes and pathways, including mTOR-sensitive translation and offsetting^46^, ISR-sensitive translation^22,51^, translation activated upon high eIF4E expression^44^, translation suppressed upon KD of eIF4A1^48^, U34-modified tRNA-dependent offsetting^49^, DHX9-dependent translation^41^, SNAT2-dependent translation^47^, high p-eIF4E-dependent translation^45^, and eIF4E2-dependent translation^50^. Additional signatures of eIF4G^43^ (GSE11011), DAP5^42^ (GSE115142), and mTOR-dependent^46^ (GSE76766) translation were obtained from supplemental material in published studies or following anota2seq analysis.

### Gene ontology enrichment analysis

Gene ontology enrichment analysis was performed using the Cytoscape (v.3.8.2.)^95^ plug-in ClueGO (v.2.5.8)^96^ with genes identified by anota2seq analysis as regulated via “translation”, “mRNA abundance”, and “buffering” in response to hypoxia treatment in T47D and H9 cell lines. The background for each cell line was defined as all genes passing expression thresholds described above for anota2seq. Enrichments were selected based on the following criteria: *FDR* cutoff = 0.05, Correction Method Used = Benjamini-Hochberg, Statistical Test Used = Enrichment (Right-sided hypergeometric test), Kappa= 0.4, Min. Percentage = 10, Min GO Level = 3, Max GO Level = 8, Number of Genes = 5, GO Fusion = false, GO Group = true, Over View Term = SmallestPValue, Group By Kapp Statistics = true, Initial Group Size = 1, Sharing Group Percentage = 50.0, Ontology Used = GO_BiologicalProcess-EBI-UniProt-GOA-ACAP, Evidence codes used = All, Identifiers used = SymbolID.

### GPF-DIA proteomics

Protein precipitation and proteolysis for C48 or DMSO-treated T47D cells were carried out as previously described^97^. Cells were lysed in 4 M Urea, 2% SDS (w/v), 1% NP-40 (w/v), 50 mM Ammonium Bicarbonate (ABC) using tip probe sonication (20 x 1 sec pulses). Protein concentrations were adjusted to 1 µg/µl prior to reduction with 5 mM (TEAB) and alkylation with 30 mM iodoacetamide (IAA) for 30 mins at room temperature in the dark. 20 µg of protein lysate were precipitated at 50% Ethanol on 100 µg magnetic beads and washed with 80% Ethanol according to SP3 protocols. Proteins were digested 1:100 with LysC for 2 hrs followed by Trypsin (1:100) overnight. Digests were collected by removing digestion buffer, collecting residual peptides with water followed by 25% acetonitrile. Collected peptides were dried by vacuum centrifugation and resuspended in 0.1% formic acid for UPLC-MS/MS analysis.

Ultraperformance liquid chromatography with tandem mass spectrometry (UPLC-MS/MS) was then performed using the gas phase fractionation data-independent acquisition (GPF-DIA) method of analysis^98^. 1 µg of lysate digest was sampled using a Neo Vanquish in trap-and-elute mode for ultraperformance liquid chromatography (UPLC). Briefly, peptide trapping and washing were performed using a PepMap™ Neo Trap Cartridge (ThermoFisher, Cat # 174500) prior to reverse-phase peptide separation using a 25 cm Aroura TS column (IonOpticks, Cat # AUR3-25075C18-TS) at 50□C and spray voltage between 1800-2000 V for 90 mins. Mobile A was composed of 0.1% Formic Acid/H_2_0 and Mobile B was composed of 0.1% Formic Acid/80% Acetronitrile/H_2_0. Peptides were loaded on the separation column at 5% Mobile B for 5 mins and increased to 10% over 5 mins, 35% over 50 mins, and 85% over 5 mins, before holding at 95% for 2 mins. For spectral library generation, 1 µg of lysate digest was serially injected to produce 100 m/z fractions across 300-1000 m/z using a staggered window scheme of 4 m/z wide windows that produce 2 m/z bins after demultiplexing, as previously described. Raw files were converted to mzML using ProteoWizard with PeakPicking = 1, Demultiplex = 10ppm, and ZeroSamples = -1. Library and sample mzML files were searched together using DIA-NN to generate a spectral library by allowing two missed cleavages and one variable modification of Oxidation. Settings for data acquisition on the Eclipse are outlined in **Supplemental File S7**.

### Analysis of GPF-DIA proteomics and RNA-seq

Paired proteomics and RNA-seq data from C48 and DMSO-treated T47D cells were independently normalized (median, and TMM normalization, respectively) and filtered to retain only genes detected in both datasets, with no missing or 0 values across all samples. Changes in protein and mRNA after C48 treatment were analyzed using the anota2seq algorithm^36^ with the normalized and filtered count data. Batch effects were accounted for in the models by including sample replicates using the “batchVec” argument in anota2seq. The threshold settings were used [minSlopeTranslation = -1, maxSlopeTranslation = 2, minSlopeBuffering = -2, maxSlopeBuffering = 1, deltaPT = deltaP = deltaTP = deltaT = 0.1] with a significance threshold maxRvmPAdj = 0.15 (corresponding to an FDR < 0.15). Genes were classified into two modes of regulation (changes in protein levels that are congruent with changes in mRNA abundance, and changes in protein levels occurring independently of mRNA levels) using the “anota2seqRegModes’ function.

### Stable isotope tracing by GC-MS

T47D cells were grown in normoxia or hypoxia (20 and 0.5% O_2_, respectively) for 24 hrs. Cells were incubated with media containing 1 mM unlabeled pyruvate for 2 hrs, then changed to 1 mM labeled ([U-13C])-pyruvate media for 15 mins (Cambridge Isotope Laboratories, MA, USA). Steady-state samples were extracted simultaneously with tracing samples. Dishes were washed in cold saline solution (9 g/L NaCl) and quenched in 80% methanol. Lysates were sonicated at 4 °C for 10 mins and spun for 10 mins (14,000 rpm, 4 °C). The internal standard (750 ng myristic acid-D27) was added to supernatants and dried overnight in a speedvac (Labconco) at 4°C. The next day, pellets were resuspended in methoxyamine hydrochloride (10 mg/ml in pyridine), sonicated and vortexed three times (30 sec each) and spun for 10 mins. Samples were heated to 70 °C for 30 mins. Derivatization with N-tert-butyldimethylsilyl-N-methyltrifluoroacetamide (MTBSTFA) was performed at 70 °C for 1 hr. Samples were injected (1 μl) into an Agilent 5975C GC/MS (Agilent Technologies, CA, USA). Analyses were performed using GC-MC (Agilent Technologies, CA, USA). Metabolite areas were normalized by the internal standard and cell number.

### Quantification and statistical analysis

Methods for quantification and statistics are described in detail in above sections, or in corresponding figure captions.

## DATA AVAILABILITY

Raw and processed RNA-seq, ChIP-seq, ATAC-seq, and nanoCAGE data have been deposited in the NCBI GEO database under accession number GSE243418. Raw proteomics data have been deposited in the PRIDE database under accession number PXD058655. Raw metabolite data have been deposited in the MetaboLights repository under accession number MTBLS12086. Other published raw datasets used in this study can be accessed at GSE11011 (eIF4G-dependent translation signature), GSE115142 (DAP5-dependent translation signature), and GSE76766 (mTOR-dependent translation signature).

## CODE AVAILABILITY

Original code produced for the study is deposited on a Code Ocean capsule and will be available upon publication. Any additional information required to reproduce the analyses reported in this study is available from the lead contact upon request.

## ACKNOWLEDGEMENTS

The authors would like to thank William Hahn, David Root, Sebastian Igelmann, and the late Jerry Pelletier for generously sharing plasmid constructs used in this work. The authors would also like to acknowledge technical help from Shannon McLaughlan and advice from Vicent Pelechano, as well as support from the National Genomics Infrastructure in Stockholm funded by Science for Life Laboratory, the Knut and Alice Wallenberg Foundation and the Swedish Research Council, and SNIC/Uppsala Multidisciplinary Center for Advanced Computational Science for assistance with massively parallel sequencing and access to the UPPMAX computational infrastructure. This project was supported by CIHR (PS-159541) and CIHR (PS-148615) awarded to LMP. Work carried out in the IT lab was funded by CIHR (PJT-451236). IT is supported by Canada Research Chair in Regulation of mRNA Translation and Metabolism. Work carried out in the OL lab was supported by grants from the Swedish Research Council (2020-01665), the Swedish Cancer Society (22 2186), the Cancer Research Funds of Radiumhemmet (211222) and the Wallenberg Academy Fellow program. DP and KW are supported by a Cancer Research Society (CRS) Next Generation of Scientists Award. PJ is supported by a Fonds de recherche du Québec (FRQ-S) Doctoral Training award.

## AUTHOR CONTRIBUTIONS

LMP, OL, and KW conceived and designed the study. BD, LL, PJ, SC, KW, JAV, TTC, DP, LM, MJ, KT, HP and RP performed experimental work. KW, BD, PJ, RP, DP, TTC, GAL LMP, OL, and IT performed analysis and interpreted the data. KS, KW, and OL developed computational methods used for the analysis of sequencing data. KW, BD, LMP, OL, and IT wrote the manuscript, with input from all authors.

---

**Figure S1.**
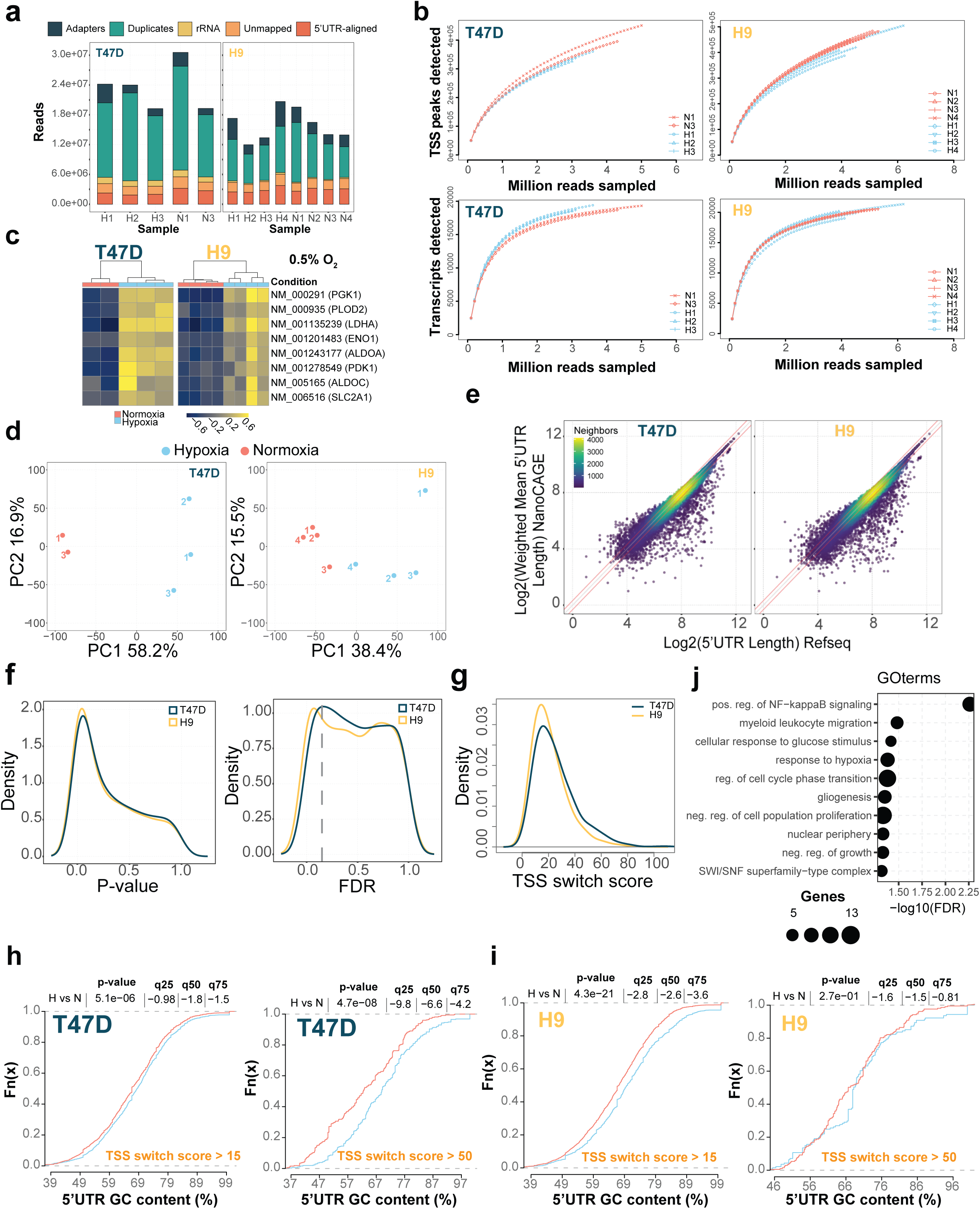
Hypoxia-induced TSS switching results in extensive remodeling of 5’UTRs. **a)** Summary of preprocessing of nanoCAGE sequencing data including filtering of RNA sequencing reads for adapters, duplicate reads, ribosomal RNA, and reads not mapping to 5’UTR regions. **b)** NanoCAGE sequencing library complexity for T47D, and H9 cells evaluated according to the number of distinct peaks (TSSs) and protein-coding transcripts (RefSeq) detected with increasing numbers of sampled sequencing reads. **c)** Heatmaps showing expression of hypoxia-responsive transcripts from nanoCAGE sequencing of T47D and H9 cells (i.e. all isoforms combined). One replicate of T47D in normoxia was discarded due to outlier characteristics. **d)** Principal component analysis of 5’UTR isoform expression in T47D, and H9 cells grown in hypoxia and normoxia quantified by nanoCAGE. **e)** Weighted mean 5’UTR length measured by nanoCAGE compared to RefSeq annotated 5’UTR length in T47D (left) and H9 cells (right). Colour gradient indicates point density. **f)** Kernel density estimation P-value (left) and FDR (right) distributions for differential TSS usage between hypoxia and normoxia in T47D, and H9 cells. Increased density of low FDRs indicates a higher frequency of changes in relative 5’UTR isoform abundance. **g)** Kernel density estimation change-point TSS switch score distributions for transcripts that undergo significant (FDR < 0.15) TSS switching under hypoxia in T47D (n = 2,552) and H9 (n = 3,423) cells. Shifts towards higher values indicate a greater difference in relative 5’UTR isoform abundance between hypoxia and normoxia. **h-i)** Empirical cumulative distribution function plots comparing the percentage of GC content between hypoxia-enriched and depleted 5’UTR sequence windows identified by change-point analysis in T47D cells. Left panel shows differences for transcripts with more modest differences in isoform abundance resulting from TSS switching (TSS switch score > 15), while right panel shows differences for transcripts with larger isoform abundance differences (TSS switch score > 50). Differences shown for T47D (**h**) and H9 (**i**) cells. **j)** Gene ontology enrichments for the set of genes corresponding to transcripts that underwent the same patterns of hypoxia-induced TSS switching in both T47D and H9 cells.

**Figure S2.**
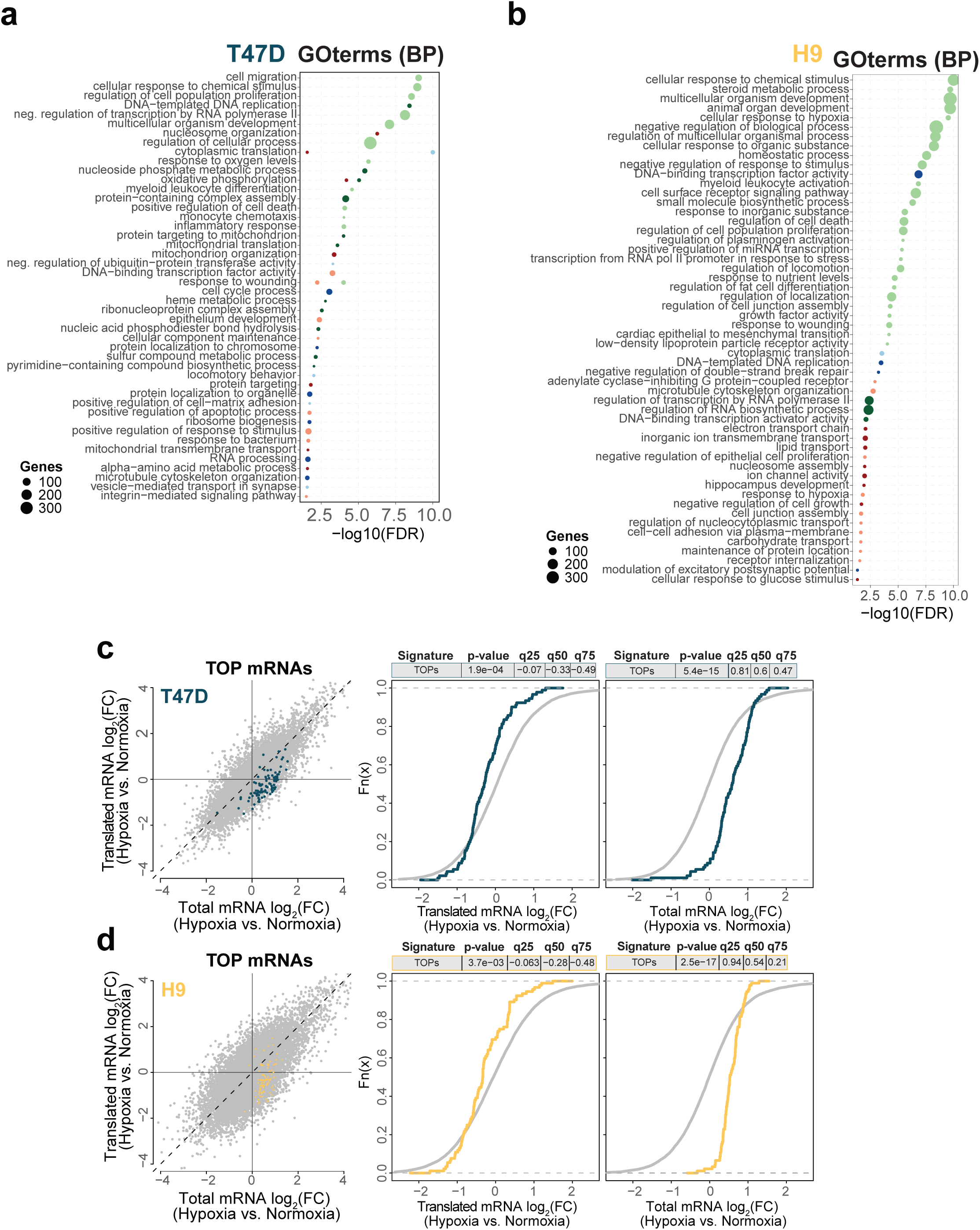
Hypoxia-induced translatome remodelling impacts many cellular processes. **a-b)** Gene Ontology enrichments for biological processes among genes assigned to each regulatory mode in Fig 2e for T47D cells (**a**) and Fig 2f for H9 cells (**b**). **c-d**) Scatterplots from anota2seq translatome analysis of T47D (**c**) and H9 cells (**d**) with the location of canonical TOP mRNAs from Philippe et al.^25^ colored (left panels). Empirical cumulative distribution functions showing the log_2_ fold changes for the TOP mRNAs in polysome-associated (middle panels), and total mRNA fractions (right panels). Grey lines correspond to the background (i.e. non-TOP genes). Significant differences between gene sets determined by Wilcoxon rank-sum test. Differences in gene expression at quantiles are indicated.

**Figure S3:**
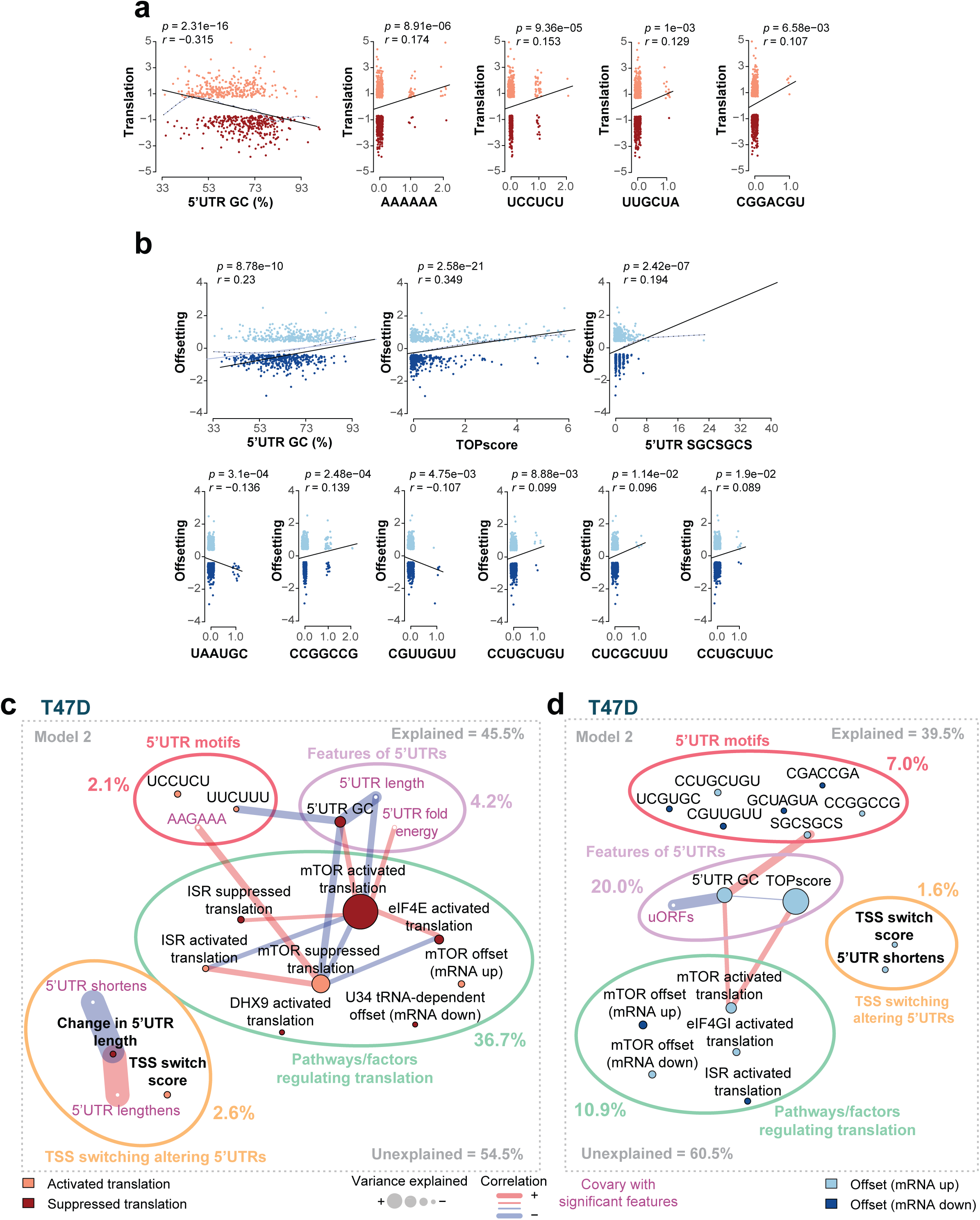
TSS switching alters regulatory 5’UTR features and shapes the hypoxia-induced translatome. **a)** Scatterplot comparing the composition of each 5’UTR feature identified in the model vs. the change in translation efficiency that occurs under hypoxia. Pearson’s correlation coefficient and p-values indicated. **b)** Same as (**a**) for translational offsetting. **c)** Networks displaying the complete results of Model 2. The total percentage of changes in translation efficiency explained and unexplained are indicated. Also indicated are the percentages of translation changes explained by different input categories. Connections between features indicate substantial correlations. Colors of the nodes indicate if the feature is associated with translation activation or suppression under hypoxia. **d)** Same analysis as **c,** but modeling changes in translational offsetting induced by hypoxia in T47D cells.

**Figure S4:**
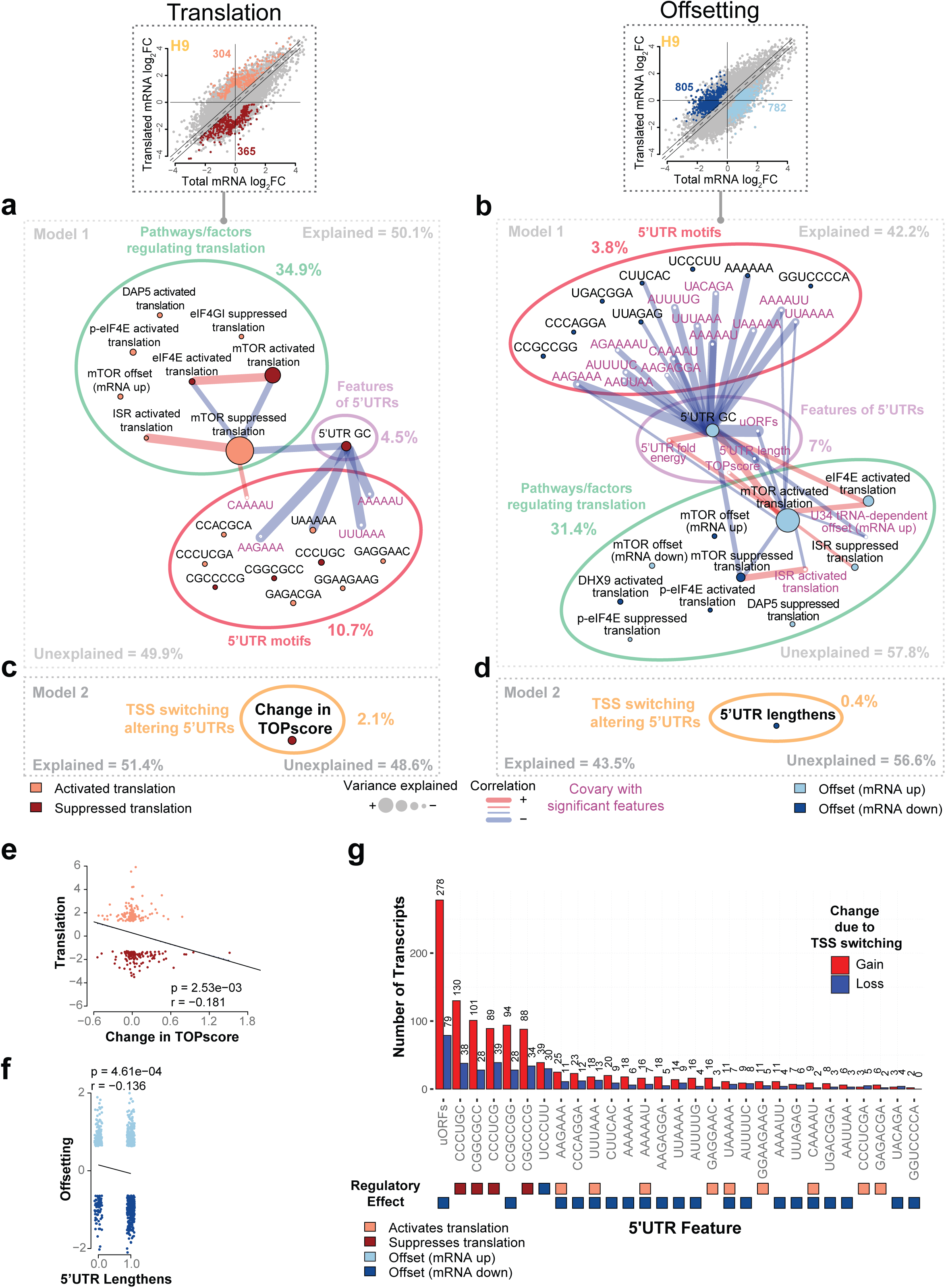
TSS switching alters regulatory 5’UTR features and shapes the hypoxia-induced translatome in H9 cells. **a)** Network plot displaying the results of Model 1. The total percentage of changes in translation efficiency explained and unexplained are indicated. Also indicated are the percentages of translation changes explained by different input categories. Connections between features indicate substantial correlations. Colors of the nodes indicate if the feature is associated with translation activation or suppression under hypoxia. **b)** Same analysis as (**a**), but modeling translational offsetting induced by hypoxia in H9 cells. **c)** Selection of the full network plot for Model 2, displaying the additional impact of adding TSS-switching signatures to Model 1 in explaining changes in translation efficiency under hypoxia in H9 cells. **d)** Selection of the full network plot for Model 2, displaying the additional impact of adding TSS-switching signatures to Model 1 in explaining changes in translational offsetting under hypoxia in H9 cells **e)** Scatterplot comparing the change in TOP score vs. the change in translation efficiency that occurs under hypoxia. Pearson’s correlation coefficient and p-values indicated. **f)** Scatterplot comparing 5’UTR lengthening events vs. translational offsetting that occurs under hypoxia. Pearson’s correlation coefficient and p-values indicated. **g)** Barplot indicating the number of transcripts that gain or lose 5’UTR elements identified in either Model 1 or 2 in H9 cells. Transcripts were considered if more than 10% of the expressed pool of 5’UTR isoforms gained or lost the 5’UTR element. The translation mode associated with each element is indicated by colored squares below.

**Figure S5:**
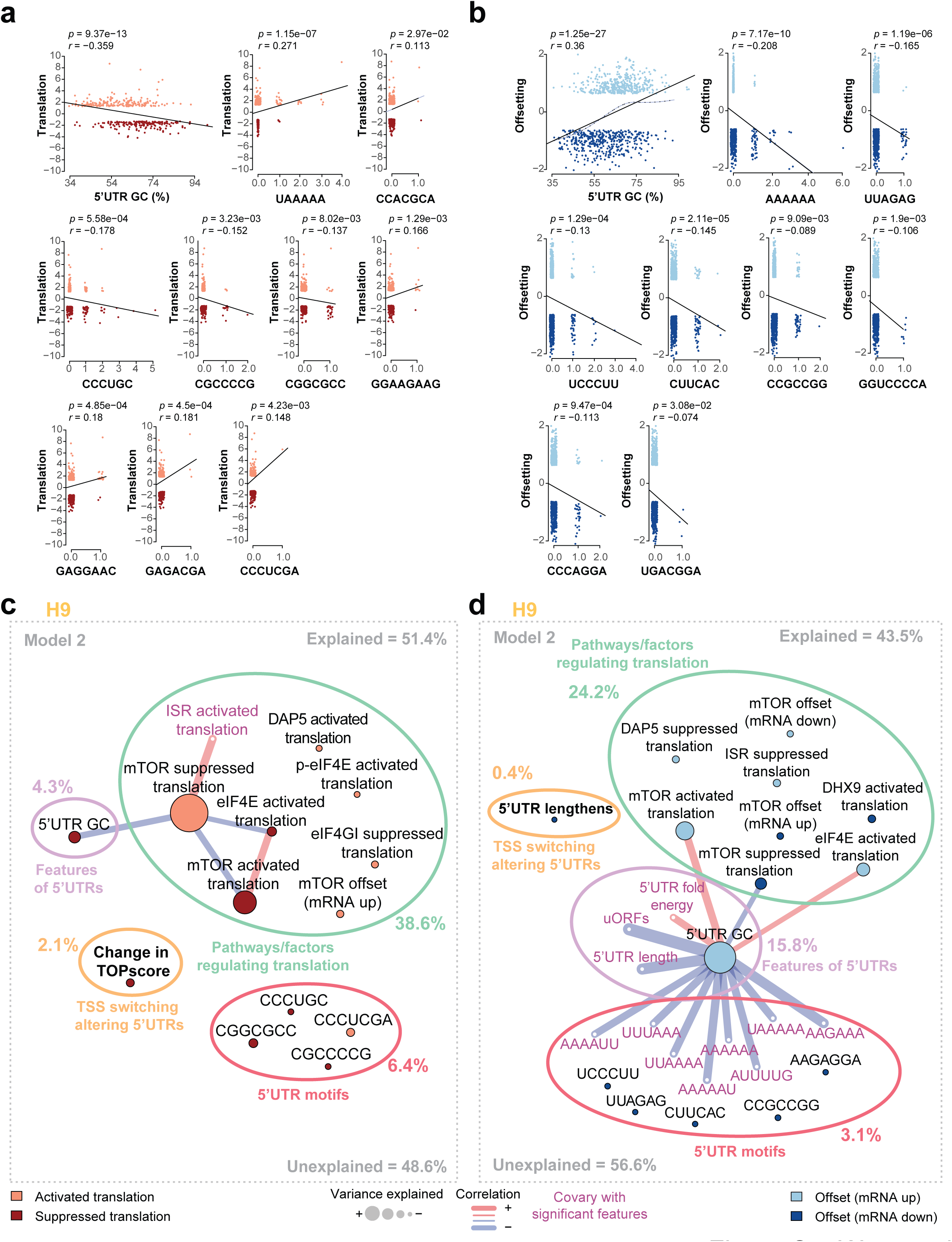
TSS switching alters regulatory 5’UTR features and shapes the hypoxia-induced translatome in H9 cells. **a)** Scatterplot comparing the composition of each 5’UTR feature identified in the model vs. the change in translation efficiency that occurs under hypoxia. Pearson’s correlation coefficient and p-values indicated. **b)** Same as (**a**) for translational offsetting. **c)** Network plot displaying the complete results of Model 2 in H9 cells. The total percentage of changes in translation efficiency explained and unexplained are indicated. Also indicated are the percentages of translation changes explained by different input categories. Connections between features indicate substantial correlations. Colors of the nodes indicate if the feature is associated with translation activation or suppression under hypoxia. **d)** Same analysis as (**c**), but modeling changes in translational offsetting induced by hypoxia in H9 cells.

**Figure S6.**
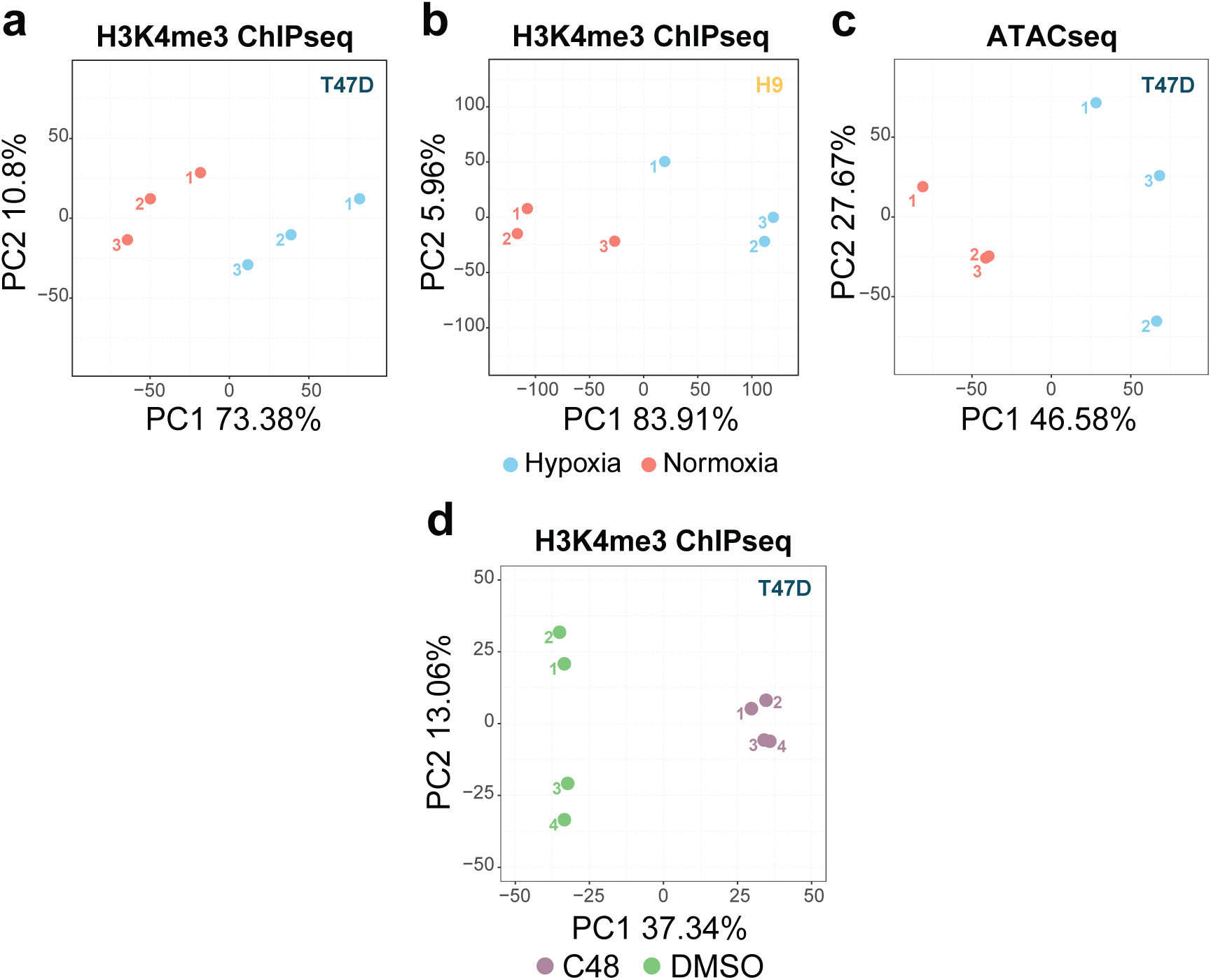
Hypoxia-induced TSS switching is associated with altered H3K4me3 and changes in nucleosome context. **a-b)** Principal component analysis of quantification of H3K4me3 peaks surrounding TSSs from T47D (**a**) and H9 (**b**) cells grown in hypoxia and normoxia and quantified by ChIPseq (n = 3). **c)** Principal component analysis of quantification of ATAC-seq peaks at TSSs from T47D cells grown in hypoxia and normoxia and quantified by ATAC-seq (n = 3). **d)** Principal component analysis of quantification of H3K4me3 peaks surrounding TSSs from T47D cells treated with C48 or DMSO for 24 hrs quantified by ChIPseq (n = 4).

**Figure S7.**
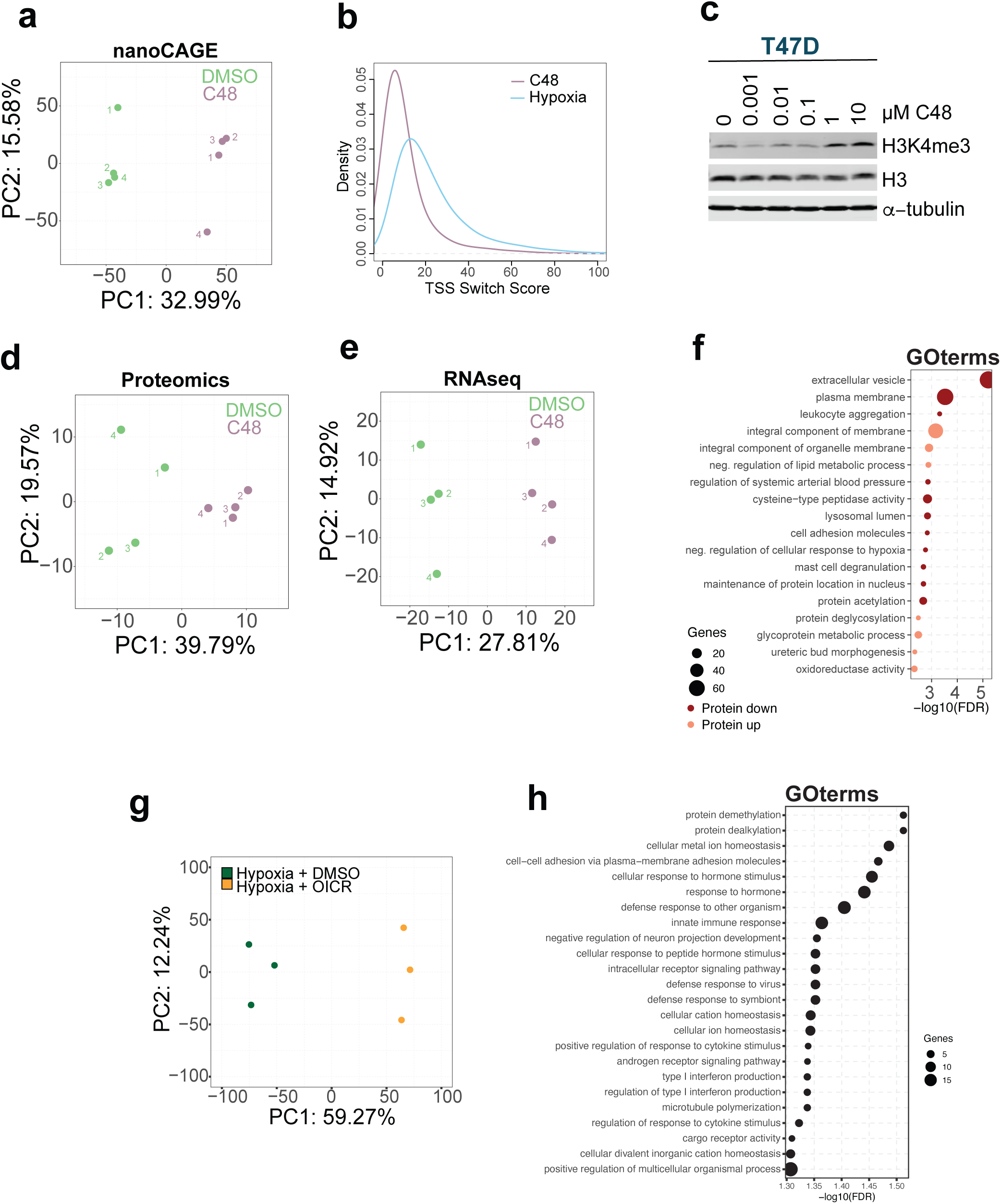
Inhibition of KDM5 induces TSS switching that remodels 5’UTRs and is associated with proteome changes. **a)** Principal component analysis of 5’UTR isoform expression measured by nanoCAGE sequencing in T47D cells treated with 10 μM C48 for 24 hrs, and DMSO-treated controls (n = 4). **b)** Kernel density estimation change-point TSS switch score distributions for transcripts that undergo significant (FDR < 0.15) TSS switching after C48 treatment (n = 3,287), compared to hypoxia treatment in T47D cells (n = 2,552). Shifts towards higher values indicate a greater degree of 5’UTR isoform expression differences. **c)** Immunoblots of H3K4me3, and H3, with -tubulin as a loading control. C48 was titrated from 0 to 10 μM in T47D cells. Lysates were harvested after 24 hrs of treatment. **d-e)** Principal component analysis for GPF-DIA proteomics (**d**), and mRNA expression measured by RNA-seq (**e**) in T47D cells treated with 10 μM C48 for 24 hrs, and DMSO-treated controls (n = 4). **f)** Gene ontology enrichments for genes where protein levels were altered independent of changes in mRNA levels in C48-treated T47D cells. **g)** Principal component analysis of 5’UTR isoform expression measured by nanoCAGE sequencing in T47D cells co-treated with hypoxia, and either 25 µM OICR-9429 or an equivalent volume of DMSO as vehicle control (n = 3). **h)** Gene ontology enrichments for genes where hypoxia-induced TSS switching was reversed by OICR-9429 treatment.

**Figure S8:**
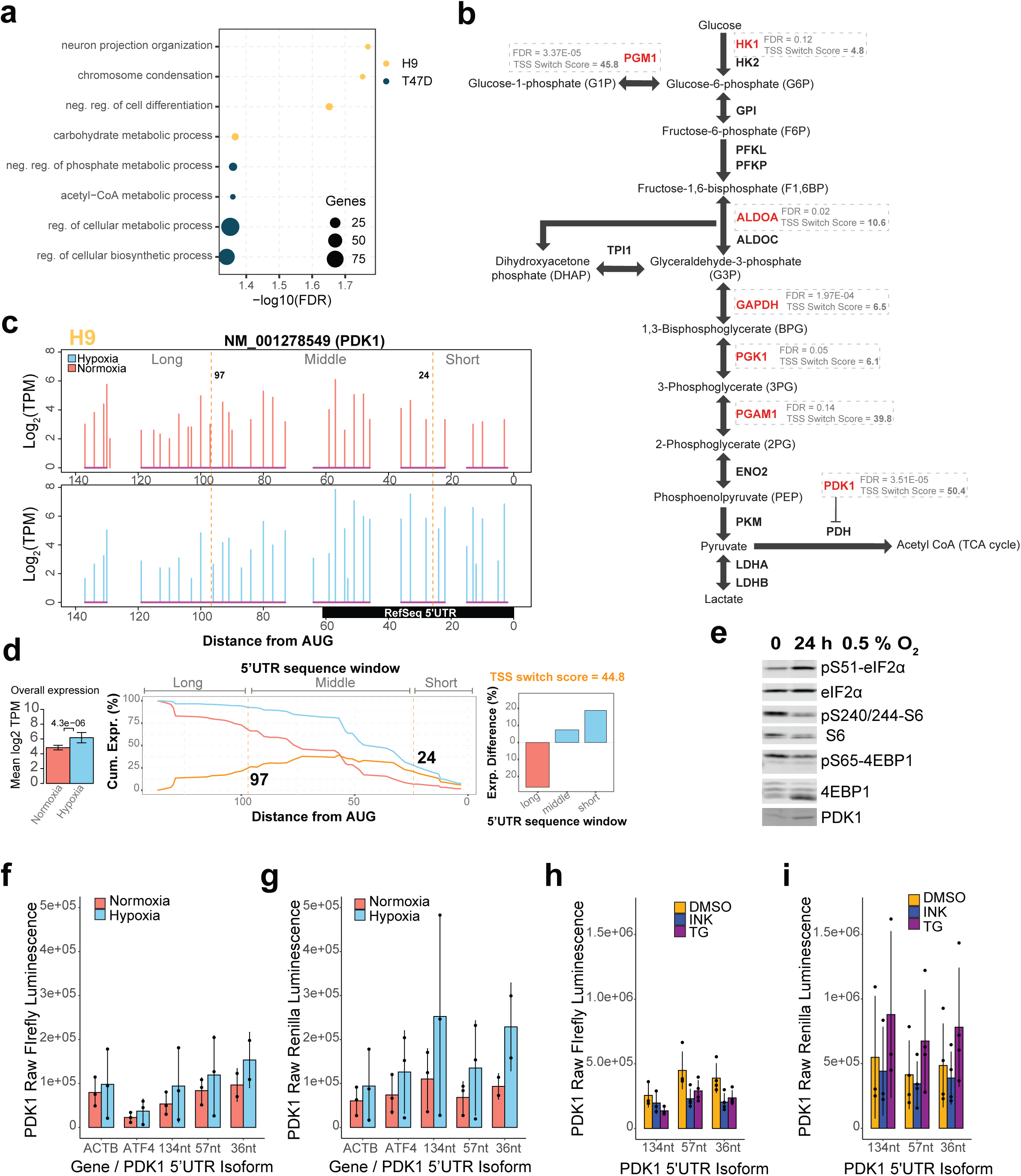
TSS switching orchestrates adaptation to hypoxia by regulating availability of differentially translated mRNA isoforms. **a)** Gene Ontology enrichments for biological processes among genes that undergo significant TSS switching under hypoxia in T47D and H9 cells. **b)** Enzymes involved in glycolysis undergo TSS switching under hypoxia. Genes with significant TSS switching events in T47D cells are marked in red. FDR values for TSS switching, and TSS switching scores denoting the degree of switching are indicated. **c)** Quantification of 5’UTR isoforms for NM_001278549 (PDK1) in hypoxia and normoxia-treated H9 cells. Same outline as Fig 1h but also indicating change-point-identified sequence windows by vertical orange lines. **d)** Change-point analysis of NM_001278549 (PDK1) identified change-points at positions 24 and 97 nucleotides upstream of the start codon (middle panel), which define shorter sequence windows with isoforms that are enriched under hypoxia and a longer window that is depleted (right panel). Difference in total mRNA expression indicated (left panel). **e)** Immunoblots of phosphorylated rpS6, 4E-BP1, and eIF2α and PDK1 from T47D cells used in polysome profiling in normoxia and hypoxia. 4E-BP1, rpS6, and eIF2α were used as loading controls. **f)** Raw Firefly luminescence values relating to Fig 7f. **g)** Raw Renilla luminescence values relating to Fig 7f. **h)** Raw Firefly luminescence values relating to Fig 7g. **i)** Raw Renilla luminescence values relating to Fig 7g.

**Figure S9:**
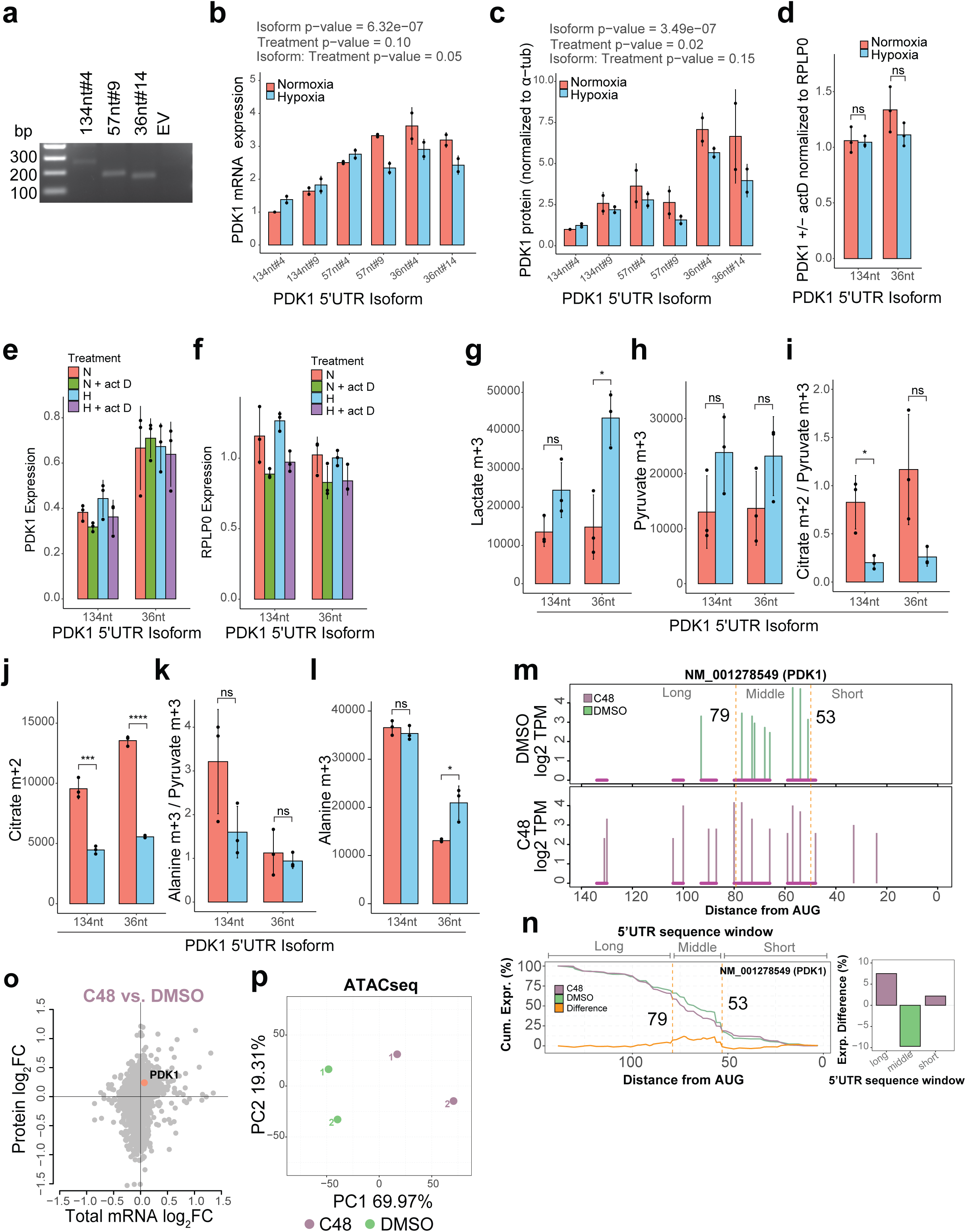
TSS switching orchestrates adaptation to hypoxia by regulating availability of differentially translated mRNA isoforms. **a)** Confirmation by PCR of the lentivirus-mediated integration of the correct isoforms in the gDNA of T47D PDK1-KO cells stably expressing 134, 57 or 36nt long 5’UTR PDK1 isoforms. PCR spans the region between the SV40 promoter and the beginning of the PDK1 ORF. **b)** Quantification of PDK1 5’UTR isoform expression in hypoxia and normoxia-treated T47D cells using RT-qPCR and normalized to 134nt#4 normoxia. P-values are provided for differences in PDK1 transcript levels between clones expressing different 5’UTR isoforms, differences depending on treatment, and the interaction between isoform:treatment from a linear model with the design Expression ∼ Isoform + Treatment + Replicate + Isoform:Treatment. Bars represent the mean and error bars indicate standard deviation (n = 2). **c)** Densitometry quantification of PDK1 protein measured by immunoblot in hypoxia and normoxia-treated T47D cells. P-values are provided as in **h**. Bars represent the mean and error bars indicate standard deviation (n = 2). **d)** Stability of PDK1 5’UTR isoforms is equal under hypoxia. T47D PDK1-KO cells stably expressing 134, or 36 nt long 5’UTR PDK1 isoforms in frame with P2A and eGFP were maintained in normoxia or hypoxia for 24 hrs and treated with actinomycin D for 16 h. The ratio of PDK1 transcripts (+/- actD) vs. RPLP0 transcripts (+/-actD) was determined by RT-qPCR (relative standard curve). Bars indicate the mean, with error bars representing the standard deviation. Significant differences determined by two-tailed t-test. * indicates p-values < 0.05. **e)** Quantification of PDK1 transcripts (+/-actD) determined by RT-qPCR (relative standard curve) used to calculate rations in (**d**). Bars indicate the mean, with error bars representing the standard deviation. **f)** Quantification of RPLP0 transcripts (+/-actD) determined by RT-qPCR (relative standard curve) used to calculate rations in (**d**). Bars indicate the mean, with error bars representing the standard deviation. **g-h)** Labelled metabolite levels of lactate m+3 (**g**), and pyruvate m+3 (**h**) from T47D cells expressing different PDK1 5’UTR isoforms grown in hypoxia or normoxia for 24 hrs. Metabolites were measured using stable isotope tracing by GC-MS. Levels are calculated as areas normalized to cell count and internal standard (myristic acid-D27). Bars represent the mean and error bars indicate standard deviation (n = 3). **i)** The ratio of labelled citrate m+2 / pyruvate m+3 from T47D cells with KO of endogenous PDK1 and re-expression of individual PDK1 5’UTR isoforms grown in hypoxia or normoxia for 24 hrs. Metabolites were determined using stable isotope tracing by GC-MS. Bars represent the mean, and error bars the standard deviation (n = 3). Significance was determined by two-tailed t-test. * indicates p-value < 0.05, ** indicates p-value < 0.005. **j)** Labelled metabolite levels of citrate m+2, as in (**g-h**). **k)** The ratio of labelled alanine m+3 / pyruvate m+3, as in (**i**). Bars represent the mean, and error bars the standard deviation (n = 3). ** indicates p-value < 0.005. **l)** Labelled metabolite levels of alanine m+3, as in (**g-h**). **m)** Quantification of 5’UTR isoforms for NM_001278549 (PDK1) in 10 µM C48 and DMSO-treated T47D cells. Change-point-identified sequence windows indicated by vertical orange lines. **n)** Change-point analysis of NM_001278549 (PDK1) identified change-points at positions 53 and 79 nucleotides upstream of the start codon (left panel), which define shorter sequence windows with isoforms that are enriched and depleted with C48 treatment (right panel). **o)** Scatter plot of protein (from GPF-DIA proteomics analysis) vs. total mRNA log_2_ fold changes in T47D cells (C48 vs. DMSO) from Fig 5k with PDK1 indicated. Anota2seq analysis identified a significant increase in protein level, independent of mRNA level. P-value < 0.05. **p)** Principal component analysis for quantification of ATAC-seq peaks at TSSs in T47D cells treated with C48 or DMSO for 24 hrs (n = 2).

